# The protein turnover and trafficking of Chlamyopsin6 is regulated by IFT88 and IFT52 in the *Chlamydomonas reinhardtii*

**DOI:** 10.64898/2025.12.11.693822

**Authors:** Kumari Sushmita, Sunita Sharma, Rajani Singh, Suneel Kateriya

## Abstract

Microbial rhodopsin-based optogenetics has been widely applied to diverse mammalian and plant cell types for controlling membrane potential mediated responses. However, trafficking of optogenetically active protein to the desired subcellular organelle is still a major concern in optogenetic field. This could be resolved by studying the trafficking mechanism of optogenetically active protein in the native system. Current study is focused on the trafficking of two of the microbial rhodopsins named Chalmyopsin5 and Chlamyopsin6 in a green alga, *Chlamydomonas reinhardtii*. Chlamyopsin5 and Chlamyopsin6 are modular in nature and possess rhodopsin, histidine kinase, response regulator and cyclase domain in tandem. Immunolocalization of Chlamyopsin5 and Chlamyopsin6 in wild strain suggests their different subcellular localization; Chlamyopsin5 in eyespot and Chlamyopsin6 in flagella. Extensive immunocytochemistry of Chlamyopsin5 and Chlamyopsin6 was performed in different intraflagellar transport (IFT) components-defective strains of *Chlamydomonas* to dissect their trafficking mode to the destined subcellular compartment. Our results indicated the trafficking of Chlamyopsin5 to the eyespot to be independent of IFT machinery while Chlamyopsin6 to the flagella to be IFT dependent. Further, we demonstrate that IFT88 and IFT52 stabilizes Chlamyopsin6 and IFT20 interacts with Chlamyopsin6 in *Chlamydomonas.* Protein interactome of Chlamyopsin5 and Chlamyopsin6 indicate their role in nitrogen assimilation, gametogenesis and photoprotection in co-ordination with other photoreceptors. Collectively, our study enabled us to understand the targeting of Chlamyopsins to the subcellular compartment (eyespot and flagella). This study is important to expand optogenetic application of microbial type modular rhodopsin with histidine kinase and response regular. Further research in this direction is required to resolve the current challenge of targeting of optogenetic protein to desired subcellular compartment.

**Graphical abstract:** 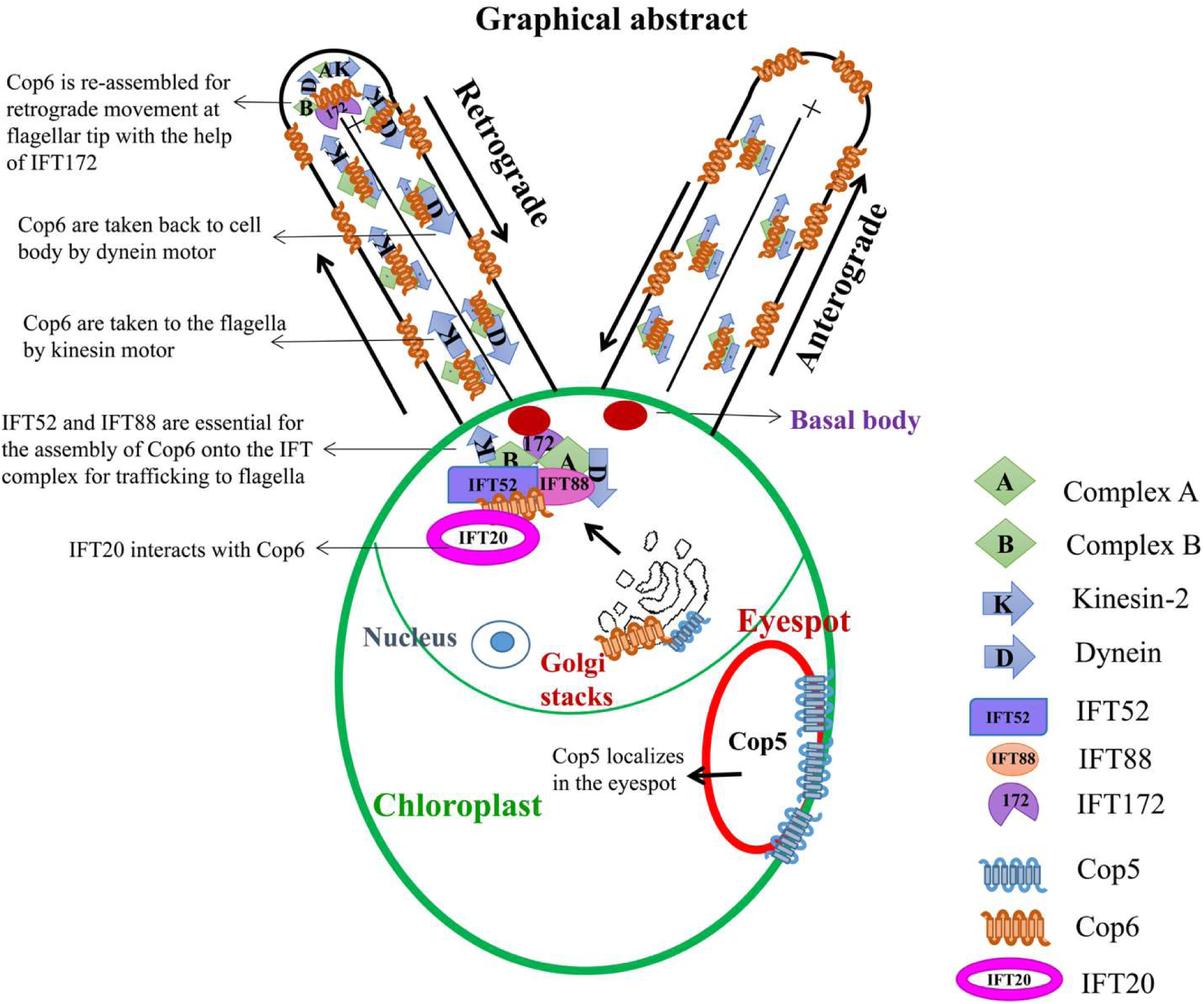

## 1. Introduction

Rhodopsins are membrane-embedded light-sensing proteins widely distributed in all the domains of life. The protein moiety in rhodopsin is bound to the light-sensing chromophore (retinal) via Schiff base linkage with the conserved lysine in the seventh helix [1]. The isomeric form of retinal in the ground state broadly divides rhodopsin into two categories- animal-type rhodopsin and microbial-type rhodopsin. Animal-type rhodopsins are further categorized as vertebrate and invertebrate rhodopsins present in photoreceptor cells in vertebrate eyes and also in the brain of invertebrates. Animal-type rhodopsins have a major role in vision and circadian clock regulation [1]. Microbial-type rhodopsins are also further categorized based on their functions as light-driven pumps, channels, sensory, or enzyme rhodopsins [1]. While optogenetic application of animal-type rhodopsin is limited due to involvement of multiple components in signaling, however, microbial-type rhodopsins are widely used for controlling the membrane potential-driven responses in diverse cell types, including mammalian and plant cells [2,3,4,5,6,7]. Despite of immense potential of rhodopsins as an optogenetic tool, the major challenges are their targeting to the required subcellular organelle or membrane surface [8,9,10]. This could be resolved to a greater extent by studying the trafficking mechanism of microbial-type rhodopsins. In this respect, trafficking of animal type rhodopsins is well studied [11]. Within the photoreceptor cells (rod and cone), rhodopsins are synthesized in the endoplasmic reticulum (ER) of the inner segment and trafficked to the Golgi complex via a COP (coatomer) mediated pathway [12]. From the Golgi complex, rhodopsins are loaded on the Golgi-derived vesicles that integrates with membrane in the vicinity of the ciliary base [11]. From the ciliary base, rhodopsins are taken by intraflagellar transport (IFT) proteins along the connecting cilium to the outer segment, and from where they get integrated into the membrane where signaling takes place [13,14].

IFT is the motor-driven bidirectional movement of proteins within the cilia, defects in which lead to ciliopathies [15,16,17]. IFT is composed of two complexes- complex A and complex B [15,16,17]. Complex A consists of 5-6 proteins along which cargoes are driven by dynein motor from the tip to the base of cilia/flagella (retrograde movement) [15,16,17]. Complex B consists of 12-16 proteins and with accompanying cargoes, is driven by kinesin motor from the base to the tip of cilia/flagella [15,16,17]. Among the IFT components, IFT52 and IFT88 are of prime importance since they form the core component of complex B required for assembly and stabilization of complex [18,19]. IFT172 of complex B mediates the switch from anterograde to retrograde movement and reassembly at the ciliary tip [20]. IFT20, one of the anterograde IFT components is unique in terms of its localization and function. IFT20 localizes at the basal body as well as Golgi complex and is associated with post-Golgi trafficking of animal type rhodopsin and other membrane proteins [11,21,22]. Though the signaling and trafficking of animal-type rhodopsin is well studied, the trafficking of microbial-type rhodopsin is lacking.

We have selected two of the microbial type rhodopsins (Chamyopsin5 and Chlamyopsin6) [23] belonging to the enzymerhodopsin category for studying the trafficking mechanism in *Chlamydomonas*. Chlamyopsin5 and Chlamyopsin6 (would be referred to as Cop5 and Cop6 hereafter) possess histidine kinase (HK), response regulator (RR), and cyclase domain in tandem and are therefore named as histidine kinase rhodopsin (HKR) or Cyclopsin (Cyclop). Biochemical characterization of purified rhodopsin domain of Cop5 (HKR1) suggests its uniqueness in terms of spectral properties. Cop5 exists in two interconvertible forms: UV and blue light absorbing state. Upon blue light exposure, it converts to the UV-absorbing state and vice versa. This interconvertible state results from the isomerization of retinal [24,25,26]. Sequence analysis of Cop5 suggests that the cyclase domain lacks a few crucial residues responsible for its function and, therefore, is suspected to interact with the functional subunit, forming a heterodimer in the native system. In-vitro characterization of full-length Cop6 in *Xenopus leavis* suggests it to be a light-inhibited guanylate cyclase with supplementation of exogenous ATP [27]. Also, the signal proceeds through HK and RR since the replacement of critical residues led to the inhibition of cyclase activity [27]. However, in the native system, the signal transduction and function might be different. We studied the trafficking and function of Cop5 and Cop6 in *Chlamydomonas reinhardtii*. Extensive immunocytochemistry of Cop5 and Cop6 was performed in wild-type and conditional mutant strains defective in IFT proteins (kinesin, dynein, IFT52, IFT88, and IFT172). These conditional mutant strains show wild-type phenotypes at permissive temperature (22°C), and mutant phenotypes (retraction or loss of flagella) at non-permissive temperature. Some of the mutant strains with disrupted genes show mutant phenotypes at permissive temperatures. Co-immunoprecipitation experiment and protein interactome prediction via web-based software were performed to get insight into the interacting partners of Cop5 and Cop6. Insight into the trafficking mechanism of microbial-type rhodopsin would help us to understand their subcellular targeting and overcome the current challenge of membrane or subcellular targeting of optogenetic proteins.

## 2. Materials and Methods

### 2.1 *Chlamydomonas* strains and culture

All strains of *Chlamydomonas* were obtained from the Chlamydomonas resource center (University of Minnesota). All wild-type strains were maintained on TAP (Tris-Acetate-Phosphate) media and mutants on TAP supplemented with 0.1g/L arginine under 14hr light and 10hr dark cycles at 22°C. All temperature-sensitive IFT mutants were grown at 22°C (permissive temperature), and experiments were carried out by incubation at 33°C (non-permissive temperature). A brief technical description of algal strains used in this study is given in table S1 [28–35].

### 2.2 Immunoblotting

Proteins were first resolved on 6 to 12% SDS-PAGE and transferred onto a nitrocellulose membrane. The protein blotted membrane was blocked with 5% skimmed milk in PBST (PBS supplemented with 0.1% Tween 20) at room temperature, incubated for 1 hr. The blocked membrane was incubated overnight with the primary antibody. The membrane was washed five times, each for five min, with PBST. After washing, the membrane was incubated with HRP-conjugated anti-goat IgG (secondary antibody) for 1hr followed by washing five times, each for five min with PBST. Membrane was developed by enhanced chemiluminescence (ECL) using solution A (100mM Tris-Cl, pH 8.5 + luminol + p-coumaric acid) and solution B (100mM Tris-Cl, pH 8.5 + hydrogen peroxide), and protein bands were captured on photographic films in the dark, according to the protocol established earlier [36,37].

### 2.3 Immunolocalization

Coverslips were washed and coated with 0.01mg/mL poly-L-lysine. *Chlamydomonas* cells were seeded on poly-L-lysine-coated coverslips and kept for 15 min at room temperature. Algal cells seeded coverslips were then treated with freshly prepared 3.7% paraformaldehyde solution in PBS for approximately 4 min. Coverslips were then dipped in ice-cold absolute ethanol solution and incubated at − 20 °C for 20 min with brief agitation. Cells were rehydrated with PBS containing 250mM NaCl, followed by four washings with PBST (PBS supplemented with 0.5% Triton X-100). The permeabilized *Chlamydomonas* cells were incubated overnight with freshly diluted primary antibody at 4°C. Then washed four times, each for 5 min with PBST, followed by incubation with FITC or Alexa Fluor conjugated secondary antibody (Invitrogen, USA) for 2 h at room temperature. After washing with PBST followed by PBS, coverslips were mounted onto the acid-washed glass slides using SlowFade® Gold antifade reagent (Molecular Probes, Invitrogen, USA) in an inverted manner [37]. Slides were visualized by the Olympus Fluoview 1000 confocal microscope, according to the protocol established earlier [36,37].

### 2.4 Co-immunoprecipitation

*Chlamydomonas* cells grown for 3-4 days with 14h light and 10hr dark cycles were harvested by centrifugation at 6000rpm for 10 min. Cells were lysed in IP-lysis buffer (RIPA) supplemented with 1mM PMSF, 1mM Sodium orthovanadate, and protease inhibitor cocktail (Sigma). Cells were fractionated by sonication, and cell debris was clarified by centrifugation at 13000rpm for 15 min. The supernatant obtained was pre-cleared by incubation with the pre-immune serum of the host organism, in which respective antibody was raised, and protein A Sepharose beads (Invitrogen) for 2h at 4°C on a gyratory shaker. One mL of pre-cleared supernatant was incubated separately with the pre-immune serum and respective antibody and kept overnight at 4°C on gyration. Protein A Sepharose beads were added to it and kept for an additional 2hr at 4°C on gyration. Beads with an antigen-antibody complex were obtained by centrifugation at low speed for 10 min. The bead pellet was washed thrice with 1X PBS. Antigen-antibody complex was eluted from the beads by resuspending the beads in 0.1M glycine, pH 2.8, equal to the volume of bead pellet, followed by centrifugation at 2000 rpm for 4 min. Supernatant was mixed with 2X Laemmli dye and heated at 65°C for 30 min. Proteins were resolved on SDS-PAGE and immunoblotted with the antibody of interest [37].

## 3. Results and Discussion

### 3.1. Spatial distribution of Chlamyopsin5 and Chlamyopsin6 differ within the *Chlamydomonas* cell

The conserved domain architecture of Chlamyopsin5 and Chlamyopsin6 is mentioned in **Figure S1**. The accession number and sequence of Cop5 and Cop6 used for domain analysis is given in Table S2. To study the localization and trafficking of Cop5 and Cop6 in *Chlamydomonas reinhardtii*, the antibodies against the peptides of the rhodopsin domains of Cop5 and Cop6 were used **(Figure S1, Appendix Figure 1)**. The protocol for the generation of antibodies is described in the appendix file supplementary protocol S1. The specificity of the antibodies was determined by immunoblotting of the purified rhodopsin domain of Cop5 and Cop6, with anti-Cop5 and anti-Cop6, respectively. The antibodies were specific towards their antigens **(Appendix Figure 2)**, and full-length Cop5 and Cop6 were detected in the cellular fractions of the *Chlamydomonas* cells **(Figure S1).** The spatial distribution of Cop5 and Cop6 differs within the *Chlamydomonas* cell. Though Cop5 possesses domains similar to Cop6, its localization was found to be restricted to the eyespot **(Figure 1)**. Cop6 behaves differently from Cop5 and localizes predominantly in flagella, with a fraction of it in the eyespot **(Figure 1)**. Dissimilar cellular destinations might be accountable for the different cellular functions of Cop5 and Cop6 within the *Chlamydomonas* cells. Since Cop6 is destined for flagella in *Chlamydomonas* cells, we sought to dissect the role of IFT machinery in the trafficking of Cop6 to the flagella.

**Figure 1:**
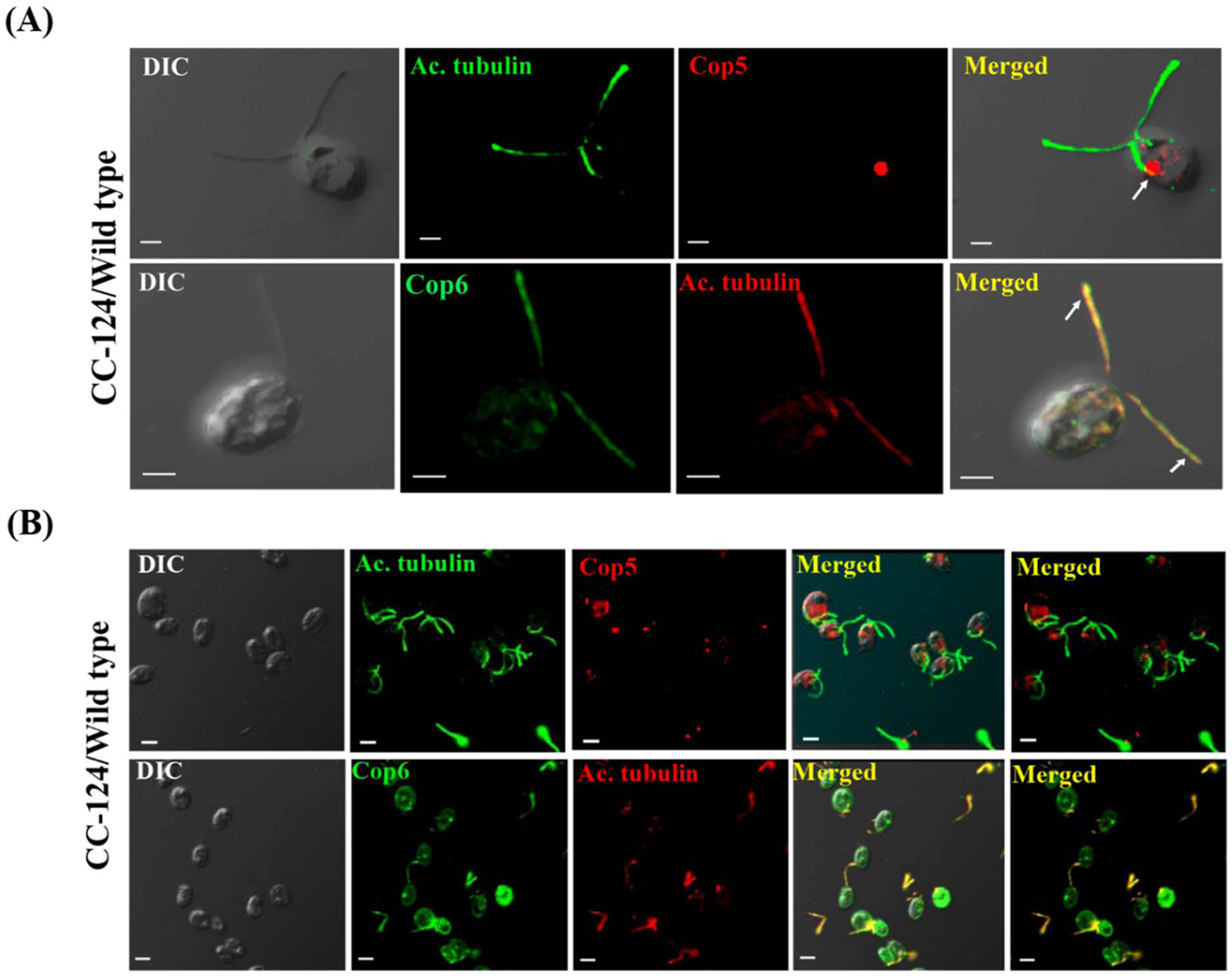
Immunolocalization of Cop5 and Cop6 in *C. reinhardtii* CC-124 (wild type) cells. (A) Represents the Cop5 and Cop6 signal in single cell. (B) Represents the Cop5 and Cop6 signal in multicells. In upper panel of A and B, immunostaining of cells was done using anti-acetylated tubulin (1:1000 dilution) (green), anti-Cop5 (1:250 dilution) (Red). The secondary antibody for acetylated tubulin and Cop5 is Alexa 488 and Alexa 546 conjugated anti-mouse IgG and anti-rabbit IgG, respectively (1:1000 dilution). In the lower panel of A and B, immunostaining of cells was done using anti-acetylated tubulin (1:1000 dilution) (Red). Fluorescence of Cop6 (1:250 dilution) (Green) was obtained with anti-Cop6. The secondary antibody for acetylated tubulin and Cop6 is Alexa 647 and Alexa 488 conjugated anti-mouse IgG and anti-goat IgG, respectively. The first panel represents DIC, and the fourth shows an overlay of the second and third panels with DIC. The experiment with wild type was performed at 22℃. Scale bar=2 μm.

### 3.2. Defects in the kinesin-2 motor protein in *Chlamydomonas* perturb the anterograde movement of Chlamyopsin6

*Chlamydomonas* Kinesin-2 is a heterotrimeric motor protein composed of two motor subunits encoded by *fla8* and *fla10*; and a non-motor subunit known as kinesin-associated protein (KAP) encoded by fla3. Kinesin-2 in *C. reinhardtii* drives the cargoes loaded IFT complex within the cilia/flagella. Previous reports elucidated the dependency of ChR1 on kinesin-2 motor protein for its trafficking to the eyespot [37]. Therefore, we investigated the involvement of kinesin-2 motor in the trafficking of Cop5 and Cop6 to the destined organelle (eyespot or flagella), via immunolocalization of Cop5 and Cop6 in a kinesin-2 motor subunit defective strains of *Chlamydomonas* (CC-1396 and CC-1919). CC-1396 and CC-1919 are a temperature-sensitive mutant strains with defects in genes encoding FLA8 and FLA10 motor subunits, respectively, affecting the assembly and transport of anterograde particles; however, retrograde movement is comparable to the wild type. The temperature-sensitive mutant strains (CC-1396 and CC-1919) have phenotypes similar to wild-type at permissive temperature (22°C), whereas at non-permissive temperature (33°C), they show characteristic features of mutations. In CC-1396 strain Cop5 localization to the eyespot is unperturbed at non-permissive temperature **(Figure S2)** while Cop6 remained concentrated near the basal body and the flagellar tip **(Figure 2, S3)**. However, even at non-permissive temperature (33°C), the Cop6 was evenly distributed along the length of flagella in wild-type cells (**Figure S4).** Thus, mislocalization of Cop6 in the absence of functional kinesin-2 indicates the dependency of Cop6 on kinesin-2 motor protein for its trafficking to flagella. On the other hand, trafficking of Cop5 to the eyespot is independent of the kinesin-2 motor. On the contrary, in CC-1919 we did not observe any change in flagellar length even at 3h time point at non-permissive temperature (33°C). No effect of flagellar length was observed at 33°C for 1h incubation while studying ChR1 localization (37), but its localization pattern was light dependent. However, no effect was observed on the localization pattern of Cop6 in CC-1919 strain at permissive and non-permissive temperatures in our study.. Earlier studies showed that 3 h is the time point at non-permissive temperature (33°C) where the average flagellar length is 50% pre-incubation length [38].

**Figure 2:**
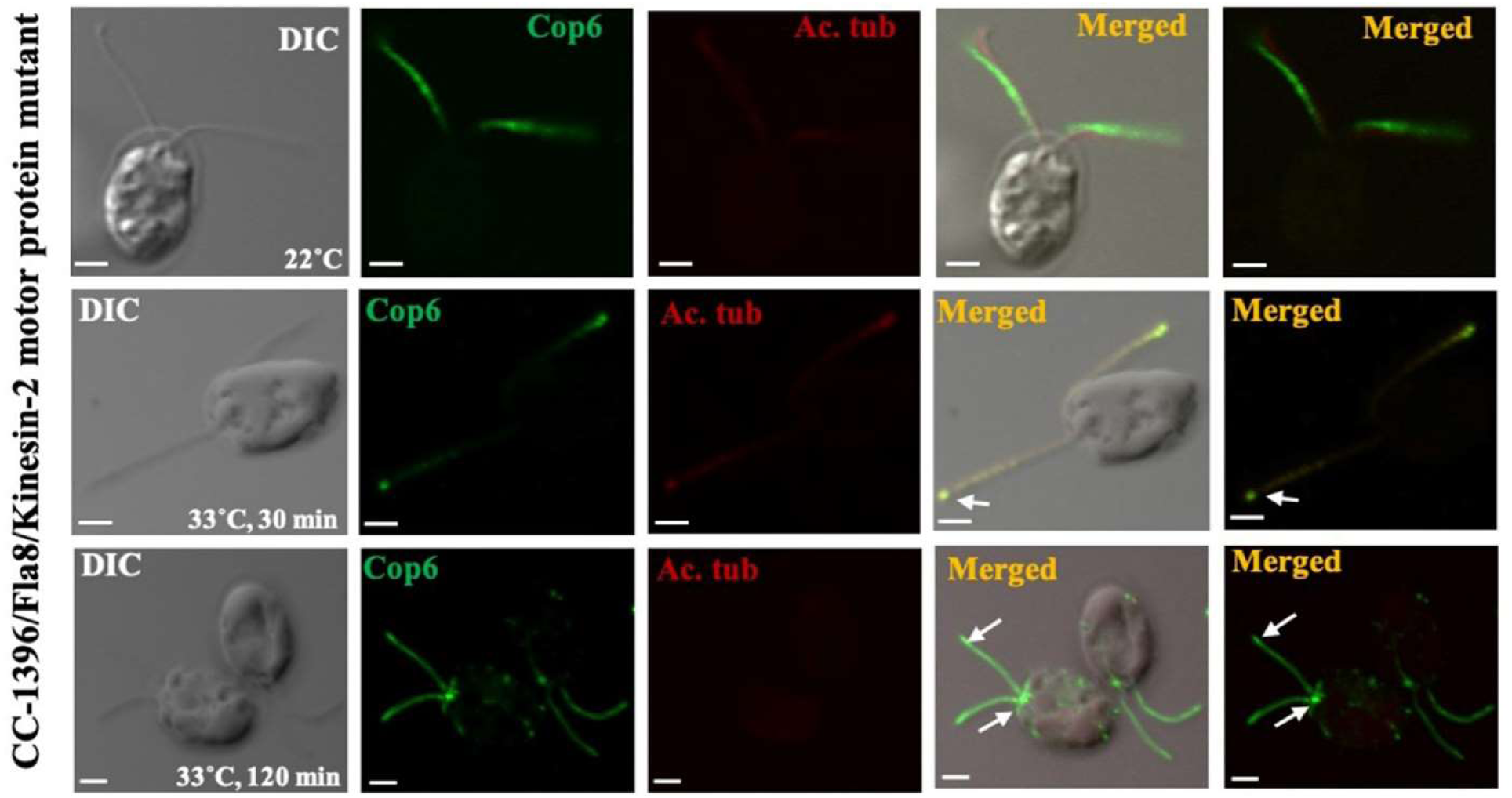
Immunolocalization of Cop6 in kinesin motor mutant strain (CC-1396) of *C. reinhardtii* cells. Immunostaining of Cop6 was done using Anti-Cop6 (1:250 dilution) (Green). The secondary antibody for Cop6 is Alexa 488-conjugated anti-goat IgG. The first channel shows DIC and the second panel depicts the fluorescence specific for Cop6, the third for acetylated tubulin (1:1000 dilution) in red. The fourth and fifth panel depicts an overlay of the second and third panels with and without DIC, respectively. The temperature for the experiment and time point is mentioned on the DIC image. Scale bar=2 μm.

### 3.3. IFT88 and IFT52 stabilize the Chlamyopsin6 in *Chlamydomonas*

The interaction of IFT52 and IFT46 via their C-terminus serves as the docking site for other IFT-B proteins, IFT70, IFT88, and the subcomplex (IFT81/74/27/25) [19]. IFT52 is crucial for the assembly of the IFT-B core complex; IFT52 mutants fail to assemble the IFT-B core complex [18]. IFT88 directly interacts with IFT52 and stabilizes the IFT-B core complex [18,19]. Thus, IFT52 and IFT88 are essential for the assembly and stabilization of anterograde IFT complex. CC-477 and CC-3943 *Chlamydomonas* strains defective in IFT52 and IFT88, respectively, were used to study the localization of Cop5 and Cop6. CC-477 (IFT52) and CC-3943 (IFT88) mutant strains are devoid of flagella even at permissive temperature (22°C). Localization of Cop5 to the eyespot is unrestrained in IFT52 and IFT88 mutants of *Chlamydomonas,* implicating that Cop5 trafficking remains unaffected by IFT disassembly and destabilization **(Figure S5)**. Surprisingly, no fluorescence for Cop6 was detected in IFT52 and IFT88 mutant strains, while in wild-type cells, fluorescence for Cop6 was detected in flagella **(Figure 3)**. In addition, a signal for acetylated tubulin was detected in both IFT52 and IFT88 mutants **(Figure 3)**. No detection of fluorescence for Cop6 in IFT52 and IFT88 led to the speculation of Cop6 degradation in the absence of IFT52 and IFT88 core components. Therefore, the protein amount of Cop6 was compared in CC-477 (IFT52 mutant) and CC-3943 (IFT88 mutant) with CC-124 (wild type) via immunoblotting. The protein amount was comparatively low in IFT52 and IFT88 mutant cell lysate than in the wild type **(Figure 4)**. The total protein amount was loaded equally in all wells. Reduced protein amount in IFT52 and IFT88 mutants with respect to wild type strengthens our speculation of Cop6 degradation due to the instability of IFT core components and the essential role of the complex for Cop6 trafficking to the destined organelle (flagella). However, further experiments would be required for the confirmation of the Cop6 degradation and the associated degradation pathway.

**Figure 3:**
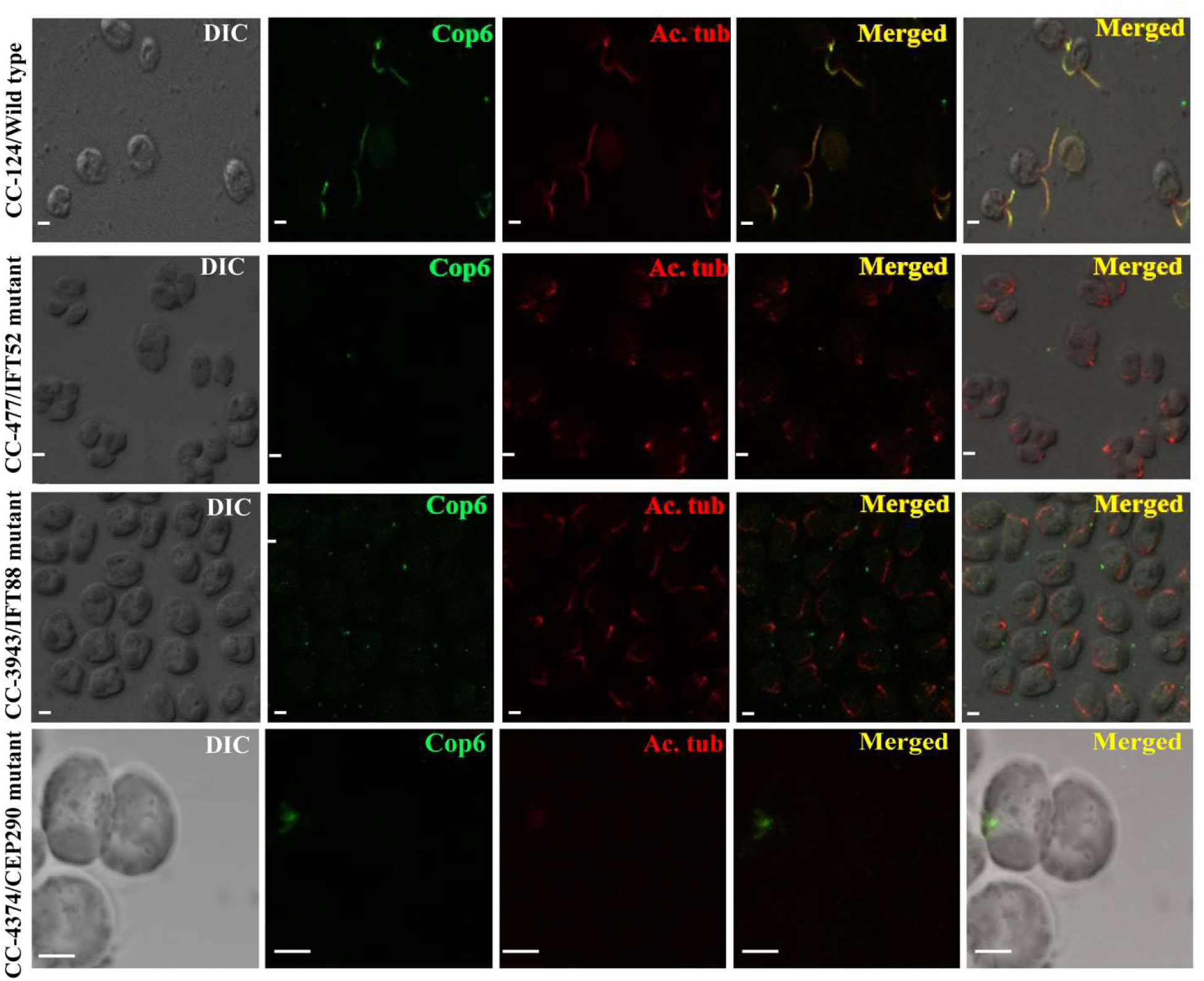
Immunolocalization of Cop6 in IFT52, IFT88, and CEP290 mutant strain of *C. reinhardtii* cells. The first panel represents DIC and the second panel is showing the signal specific to Cop6 with the primary antibody anti-Cop6 at 1:250 dilution in green. Secondary labelling was done with Alexa 488 conjugated anti-goat IgG. The third panel represents the signal for acetylated tubulin (1:1000 dilution) in red; secondary labelling was done with Alexa 647 conjugated anti-mouse IgG. The fourth and fifth panel represents the overlay of the second and third panels without and with DIC, respectively. As these are not temperature sensitive mutants, the temperature and time point is same as wild-type ( 22℃). Scale bar=2 μm.

**Figure 4:**
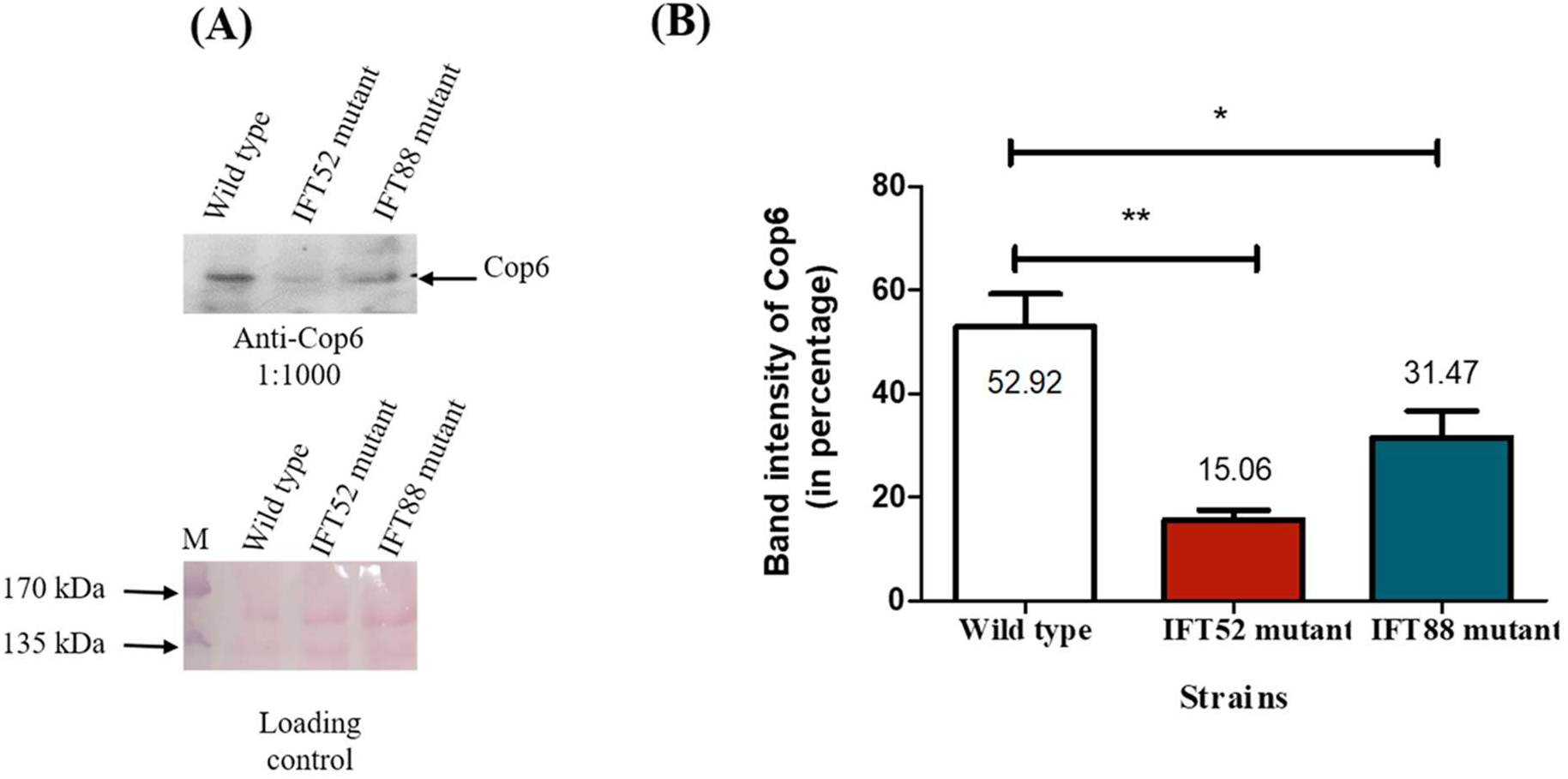
Comparison of the expression profile of Cop6 in wild type versus IFT52 and IFT88 mutants. (A) Immunoblot of cell lysate of the wild-type strain, IFT52, and the IFT88 mutant strain with anti-Cop6. (B) Represents the relative band intensity of Cop6 in wild-type strain, IFT52, and IFT88 mutant strain. Value represents mean of n=3 ± SEM. Statistical significance was assessed using unpaired t-tests; p < 0.05 was considered significant.*P < 0.05, **P < 0.001.

Taken together, we deduce that IFT core components IFT52 and IFT88 stabilize their cargo protein Cop6 and are crucial for its trafficking to flagella, whereas Cop5 stabilization and trafficking to the eyespot is independent of IFT52 and IFT88.

Further, we have performed the immunolocalization of Cop6 in CC-4374, a mutant strain of *C. reinhardtii* that lacks most of the CEP290 gene due to deletion. These mutants either lack flagella or bear very short/stumpy flagella at permissive temperature (22°C). CEP290 is a large coiled-coil centrosomal protein also shown to be present at the transition zone. The transition zone is a connecting link between the basal body and protruding cilia/flagella, acting as a gate for entry and exit for all ciliary proteins [39,40]. Disruption of CEP290 in *C. reinhardtii* leads to an imbalance of IFT A and IFT B proteins transiting the flagella at the transition zone, affecting the trafficking of ciliary proteins. Cop6 was found to be mislocalized in the cell body of CC-4374, and no signal was observed near the basal body **(Figure 3)**. Mislocalization of Cop6 suggests its impaired trafficking due to the imbalance of IFT proteins regulated through the CEP290 protein.

### 3.4. IFT172 regulates the transition of Chlamyopsin6 from anterograde to retrograde movement at the flagellar tip

IFT172 is a component of the anterograde IFT complex encoded by *fla11* [20]. IFT172 mediates the switch from anterograde to retrograde movement at the flagellar tip in *Chlamydomonas* [20]. A temperature-sensitive, flagellar assembly mutant strain of IFT172 (CC-1920) was used to study the localization of Cop6 to dissect the involvement of IFT172 in the turnover of Cop6 at the flagellar tip. CC-1920 is a temperature-sensitive, flagellar assembly mutant of IFT172 encoded by *fla11* with replaced Leucine with Proline (L_1615_P) at position 1615. The mutant strain shows wild-type-like phenotypes at 22°C, but the flagella get reabsorbed with time when incubated at 33°C. Cop6 sticks at the flagellar tip in CC-1920 strain at incubation of 30, 60, and 90 min at a non-permissive temperature (33°C) while evenly distributed within the flagella at a permissive temperature (22°C) **(Figure 5A)**. The quantification of Cop6 signal along the length of flagella was performed using ImageJ (**Figure 5B**). Accumulation of Cop6 at the flagellar tip indicates the role of IFT172 in turnover at the flagellar tip (switch of anterograde to retrograde movement).

**Figure 5:**
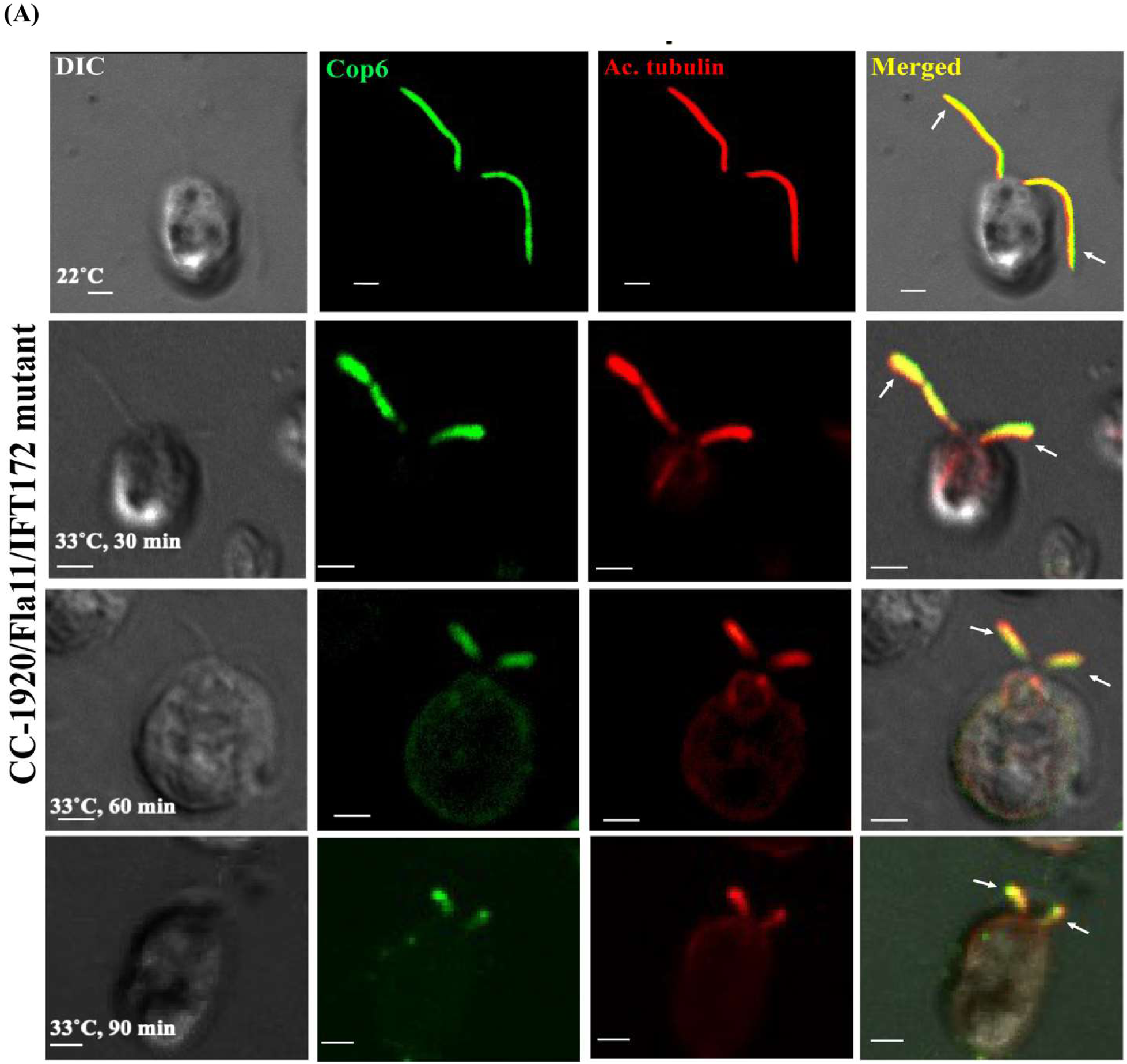
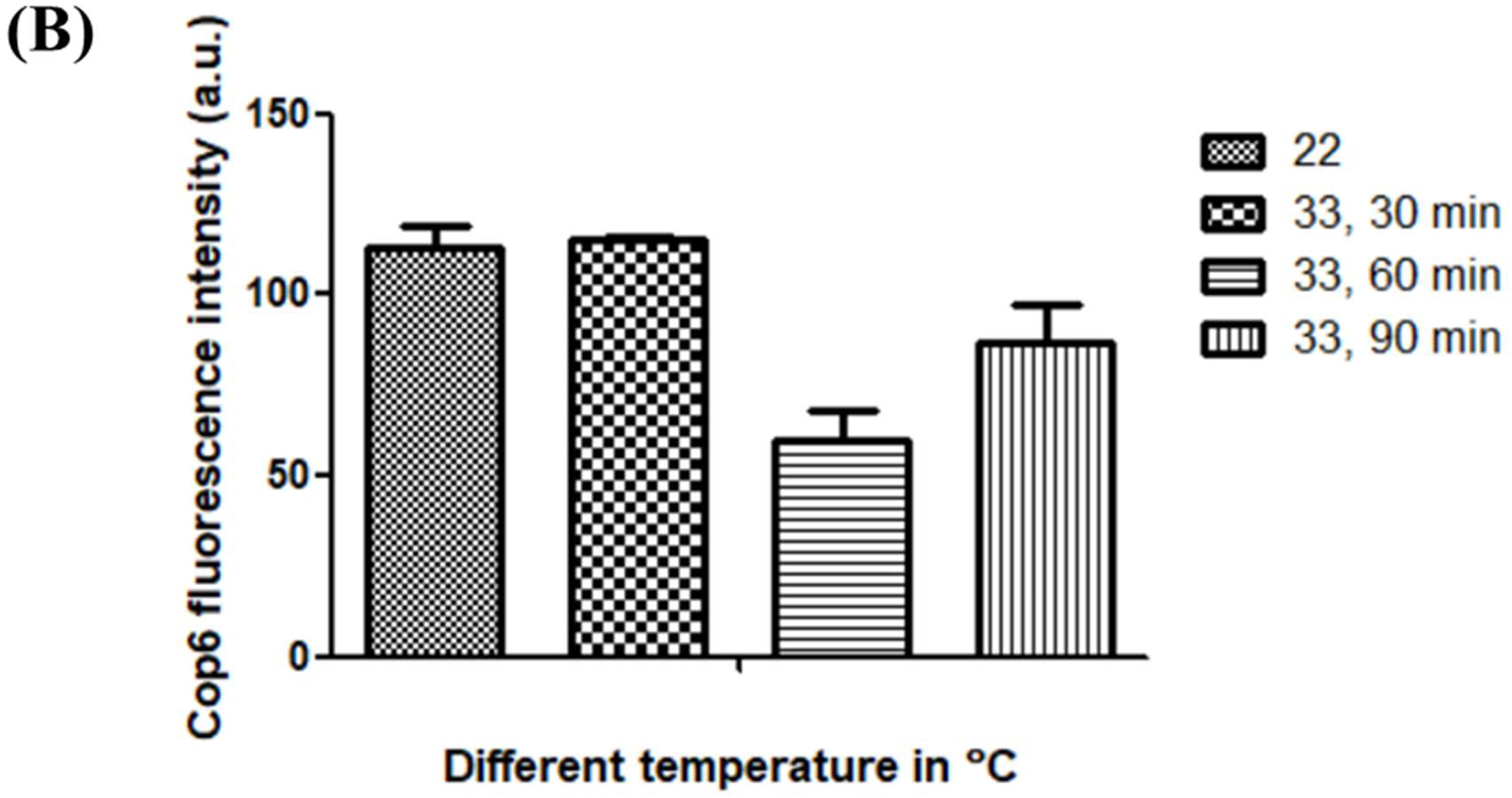
Immunolocalization of Cop6 in IFT172 mutant strain (CC-1920) of *C. reinhardtii* cells. (A) Immunostaining of cells was performed using anti-acetylated tubulin (Red) and anti-Cop6 (Green). The secondary antibody for acetylated tubulin and Cop6 was Alexa 647 and Alexa 488 conjugated anti-mouse IgG and anti-goat IgG, respectively. The first channel shows DIC and the second panel depicts the fluorescence specific for Cop6, the third for acetylated tubulin. The fourth panel depicts an overlay of the second and third panels with DIC. The temperature for the experiment and time point is mentioned on the DIC image. Scale bar=2 μm. (B) Quantification of Cop6 signal along the length of flagella using Image J at 22℃ and different time point at 33℃

### 3.5. Chlamyopsin6 interacts with IFT20 in *Chlamydomonas*

IFT20 is a part of complex B (anterograde IFT components) and is unique among all IFT components regarding its distribution within the cell. All IFT components show their presence at the basal body, while IFT20 was found associated with the Golgi complex as well as the basal body. IFT20 mediates the trafficking of ATR from the Golgi to the basal body and within the cilia/flagella [21,22]. It interacts with Rabin8, a GEF (guanosine exchange factor) for GTPase (Rab8) involved in post-Golgi trafficking of ATR in vertebrates [11]. To dissect its role in the trafficking of modular sensory-type rhodopsin (Cop5 and Cop6) in *Chlamydomonas,* the antibody was raised against the full-length recombinant IFT20 (Appendix file Supplementary protocol S2). Specificity and sensitivity of anti-IFT20 was confirmed by immunoblotting varying concentrations of recombinant IFT20 protein **(Appendix Figure 3)**. Anti-IFT20 was able to detect 50ng of recombinant IFT20. After confirmation of the specificity of the IFT20 antibody, it was used to detect the full-length IFT20 from the cell lysate of *Chlamydomonas.* Anti-IFT20 was able to detect the specific full-length IFT20 from the total cell lysate of *Chlamydomonas,* and no band was detected with pre-immune serum **(Figure S6A)**. IFT20 was seen to be localized in the flagella of *Chlamydomonas* cells **(Figure S6B)**.

Interactome prediction of IFT20 in *Chlamydomonas* suggests its interaction with other IFT components and, therefore, is an important component of IFT machinery. In the mammalian system, IFT20 was reported to form a link between IFT components and kinesin-2 by interacting with the motor protein (KIF3B) of kinesin-2 motor [41][30]. To assess the role of IFT20 in the trafficking of Cop6, the interaction of IFT20 and Cop6 was checked via co-immunoprecipitation. Anti-IFT20 was used as bait to precipitate Cop6 from the total cell lysate of *Chlamydomonas* cells. Cop6 precipitated with anti-IFT20, suggesting the interaction between IFT20 and Cop6 in the cellular system **(Figure 6)**. The interaction of IFT20 with Cop6 indicates that IFT20 might assist in the trafficking of Cop6. However, the assistance of IFT20 in trafficking Cop6 from the Golgi complex or the basal body remains unexplored.

**Figure 6:**
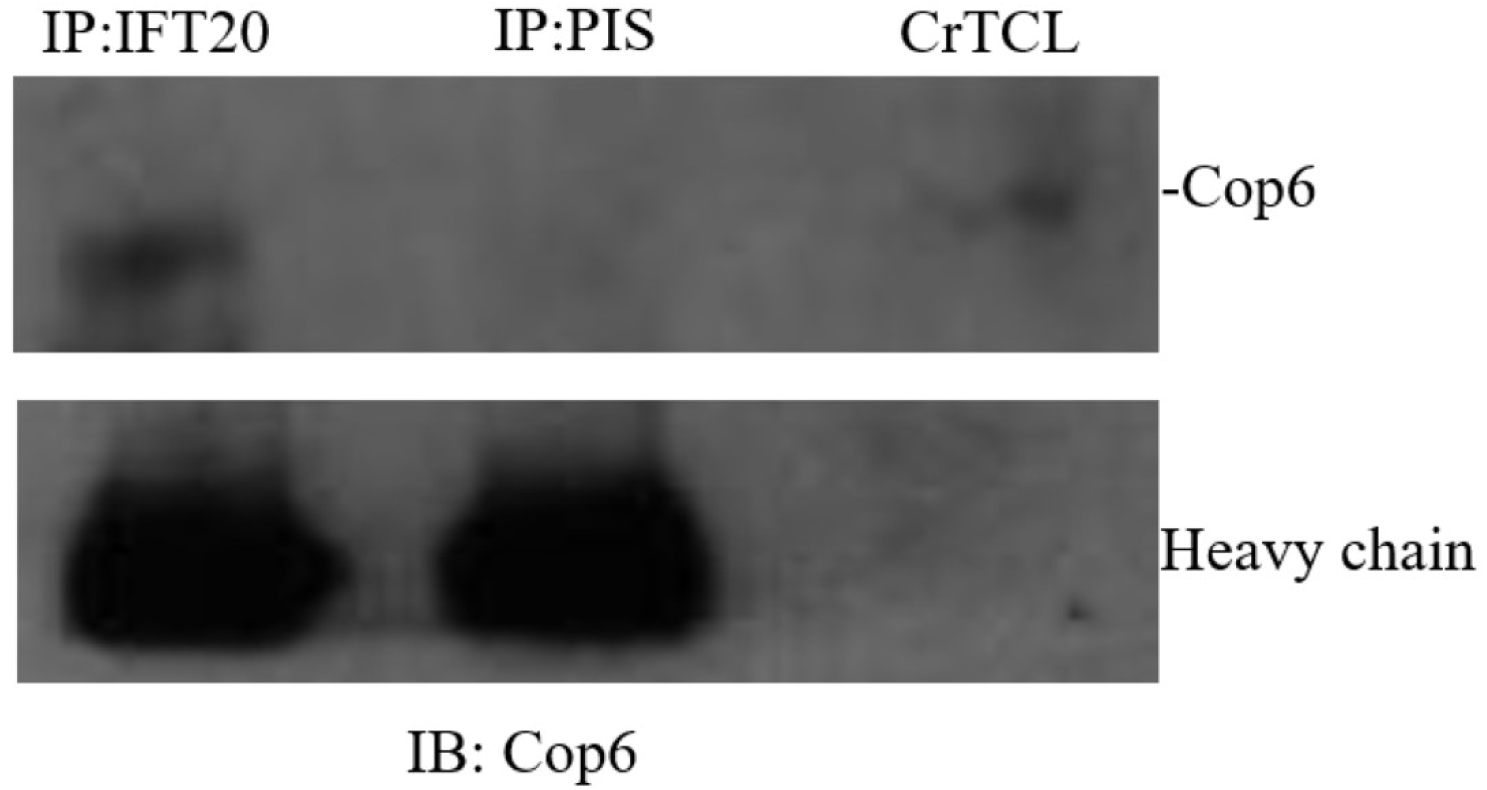
Co-immunoprecipitation of Cop6 with IFT20 antibody. Co-immunoprecipitation of Cop6 with IFT20. Anti-IFT20 (IP: IFT20) and pre-immune serum (IP: PIS) of a goat in which antibody against IFT20 was raised were incubated with the cell lysate of *Chlamydomonas*, followed by an immunoprecipitation protocol. Immunoblotting was performed with anti-Cop6. The band at the size of 250 kDa was detected in IP with anti-IFT20. The band at the corresponding position was also observed in the cell lysate utilized for IP. No band was detected in IP with pre-immune serum, reflecting the specificity of the band in IP with anti-IFT20. An equal amount of heavy chain in IP with anti-IFT20 and pre-immune serum reflects the loading control.

### 3.6. Dynein motor drives the retrograde movement of Chlamyopsin6 in Chlamydomonas

The dynein motor is composed of heavy, light, and intermediate chains. Dynein motors drive the retrograde movement of IFT components and their cargo proteins. However, some proteins return to the cell body via diffusion, including kinesin-2 [42]. To explore the retrograde movement of Cop6 via the dynein motor, *Chlamydomonas* strain (CC-4424) with a defect in the kinesin motor and dynein heavy chain was used to study immunolocalization. This mutation affects the retrograde movement of cargo proteins at a non-permissive temperature (33°C), resulting in the accumulation of cargo proteins at the tip of cilia/flagella. Accumulation of Cop6 at the flagellar tip in dynein-defective *Chlamydomonas* mutants implicates that Cop6 requires the dynein motor for its movement back to the cell body from the flagella **(Figure 7)**. Localization of Cop5 in the CC-4424 mutant strain was unperturbed at non-permissive temperature **(Figure S7)**; it localized to the eyespot, indicating the retrograde movement of Cop5 is independent of dynein motors.

**Figure 7:**
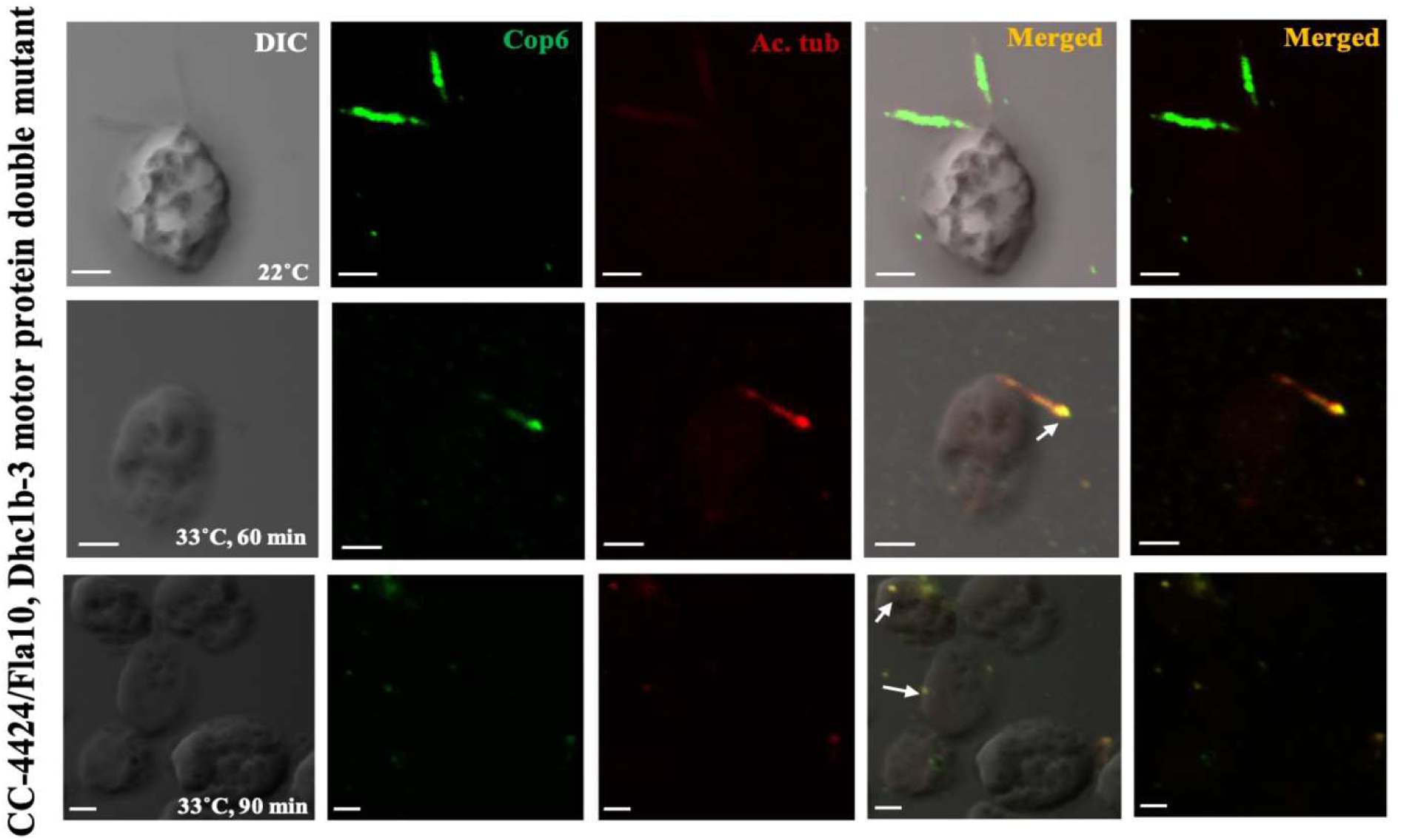
Immunolocalization of Cop6 in kinesin and dynein heavy chain motor double mutant strain (CC-4424) of *C. reinhardtii* cells. Immunostaining of cells was performed using anti-acetylated tubulin (Red) and anti-Cop6 (Green). The secondary antibody for acetylated tubulin and Cop6 is Alexa 647 and Alexa 488 conjugated anti-mouse IgG and anti-goat IgG, respectively. The first channel shows DIC and the second panel depicts the fluorescence specific for Cop6, the third for acetylated tubulin. The fourth and fifth panel depicts an overlay of the second and third panels with and without DIC, respectively. The temperature for the experiment and time point is mentioned on the DIC image. Scale bar=2 μm.

### 3.7. Speculation of Chlamyopsin5 and Chlamyopsin6 function via protein-protein interaction

To gain insight into the pathways associated with Cop5 and Cop6 and their physiological role in *Chlamydomonas*, the interacting partners of Cop5 and Cop6 were predicted through the STRING database [43]. 14-3-3 is a regulatory protein involved in diverse intracellular processes through its interactions with phosphorylated and non-phosphorylated proteins via different motifs [44]. Various canonical and non-canonical motifs have been identified for the binding of 14-3-3 to the target proteins [44,45,46,47,48].The addition of the sequence of regulatory protein (14-3-3) to the interactome established the link between 14-3-3 and phototropin [36,49,50] and, in turn, with other photoreceptors- ChR1 (Cop3), ChR2 (Cop4), UVR3 and cryptochrome (CPH1 [36,51,52,53], suggesting the complex cross-talk among photoreceptors that might regulate signaling pathways like-photosynthesis, photoprotection, eyespot development, sexual cycle or circadian rhythm. 14- 3-3 interaction with IFT complex via Actin (IDA5) suggests its probable role in trafficking. Therefore, a combined interactome was constructed using information from STRING, the literature, nano-LC-MS/MS lab data, and sequence analysis to identify a 14-3-3 binding motif **(Figure 8)**. Sequence analysis of IFT proteins and photoreceptors showed the presence of putative non-canonical 14-3-3 binding motifs (RX_1-2_pSX_2-3_S) in Phototropin, Cop5, Cop6, IFT172 and Kinesin-2 motor protein (FLA8) [28]; in addition, Dynein heavy chain 1b possesses a mode III binding motif (pS/pTX_1-2_-COOH) in the form of FLSV represent by red dotted line in the interactome. The prediction of putative binding motifs for 14-3-3 in photoreceptors and IFT proteins served as an intermediate to connect the signaling pathways. However, experimental evidence is required to prove these interactions. Cop5 in association with NIT1 (nitrate reductase), might play a role in nitrogen assimilation. The interactome of Cop6 is unknown, and therefore, no interaction was observed for Cop6. Our co-immunoprecipitation experiment suggests the interaction of Cop6 with IFT20, the nature of the interaction (direct or indirect) cannot be specified. The expression pattern of photoreceptors (ChR1, ChR2, Phototropin, LOV-HK1, LOV-HK2) and 14-3-3 genes during different phases of gametogenesis suggests their role in the sexual cycle of *Chlamydomonas reinhardtii* [36]. Cop5 and Cop6, possessing cyclase domains at their C-terminus, might have some role during gametogenesis through crosstalk with other photoreceptors and signaling protein 14-3-3. Further, cytohubba analysis was also performed to identify the hub protein governing the network **(Figure S8, Table S3).**

**Figure 8:**
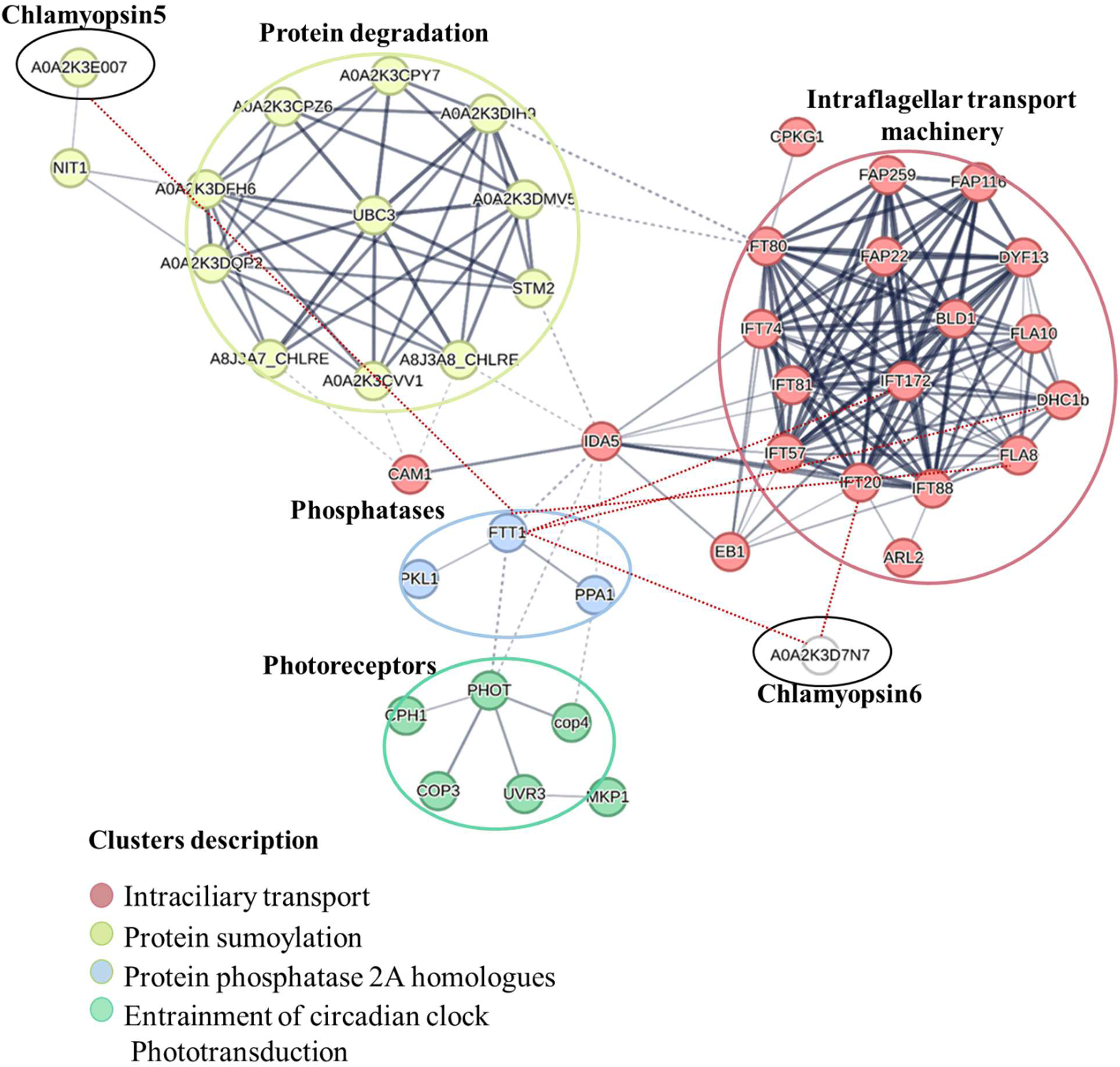
Predicted interactome of Cop5, Cop6, and 14-3-3. A combined interacting network was constructed based on the interactions reported in literature, interactions derived from the STRING database, proteins possessing a putative interacting motif for 14-3-3, and proteins detected in Nano-LC-MS/MS of lab data. The red dotted lines denote interactions based on co-immunoprecipitation and interaction prediction based on a putative interacting motif for 14-3-3. The clusters marked with different colours represent the proteins involved in particular biological processes in the organism.

## 4. Conclusions

In this study, we intended to study the IFT-mediated trafficking of two of the modular rhodopsins (Chlamyopsin5 and Chlamyopsin6) from *Chlamydomonas reinhardtii*. Despite being similar in domain organization, the spatial distribution of Cop5 and Cop6 in wild wild-type strain of *Chlamydomonas* differs. Cop5 localizes in the eyespot, whereas Cop6 localizes in the flagella of the *Chlamydomonas* cells. Further, an *in vivo* localization study of Cop5 and Cop6 in temperature-sensitive IFT motor (kinesin and dynein) mutants of *Chlamydomonas* suggested that Cop6 is dependent on IFT motors for its transport to the flagella, while transport of Cop5 to the eyespot is independent of IFT motors. The combined result of immunofluorescence and immunoblotting of Cop6 in IFT88 and IFT52 mutant strains suggested the dependency of Cop6 on IFT88 and IFT52 for stability and trafficking to flagella in the *Chlamydomonas* cells. IFT172 mediates the reassembly of Cop6 for retrograde movement at the flagellar tip. Cop6 co-precipitated with IFT20 and might serve as a link between Cop6 and IFT proteins. Protein-protein interaction suggested the role of Cop5 and Cop6 in nitrogen assimilation, gametogenesis, phototaxis/photophobic response, and photoprotection via direct or indirect interaction with other photoreceptors (ChR1, ChR2, and phototropin), and 14-3-3.Interactions should be further confirmed by pull-down assays. IFT-mediated trafficking and functional analysis of Cop5 and Cop6 would enable us to understand the selectivity mechanism of IFT machinery for microbial-type rhodopsin and their subcellular targeting. In addition to enhancing the optogenetic applicability of rhodopsin, such a study would also enable us to understand ciliopathies. In conclusion, our study indicates that the trafficking mechanisms of Cop5 and Cop6 differ in the native system. Some of the IFT components (e.g-IFT52 and IFT88) not only mediate the trafficking of Cop6 but also help in its stabilization. IFT20, directing the movement of animal photoreceptor from the Golgi complex, also interacts with Cop6. Protein interactome of Cop5 and Cop6 suggests the crosstalk among the photoreceptors and would enable us to establish its link with many light-mediated biological pathways in *Chlamydomonas reinhardtii*.

## Acknowledgement

SK is thankful to SERB ANRF (CRG/2021/00315), (EEQ/2023/000398) and DBT (BT/PR53866/BSA/33/107/2024) for providing financial support for the research project. Dr. Peeyush Ranjan and Dr. Mayanka Awasthi are acknowledged for the gift of Cop5 and Cop6 antibodies. DBT and CSIR are highly acknowledged for providing financial assistance to KS and SS, respectively. RS acknowledges the DBT-RA program (DBT-RA/2022/July/N/2560) for financial assistance.

## Author Contributions

S.K. conceived the project. K.S. designed experiments under the guidance of S.K. and drafted the manuscript with contributions from S.K., S.S., and R.S. K.S. conducted immunofluorescence, immunoblotting, Co-immunoprecipitation experiments, and PPI interactome. S.S. contributed to designing experiments, immunoblotting, immunofluorescence, PPI interactome analysis, and image processing. R.S. contributed to immunoblotting, PPI network prediction, and image processing. All authors reviewed, revised, and approved the final manuscript.

## Data availability

Data will be made available on request.

## Glossary

**Anterograde:** Directional transport from flagellar base to distal tip via kinesin-2 motors in IFT complexes.

**Retrograde:** Directional transport from flagellar distal tip base to tip via dynein motor protein in IFT complexes.

**Chlamyopsin:** Rhodopsin-like photoreceptor in *Chlamydomonas reinhardtii* mediating light responses in cell.

**Intraflagellar transport:** highly organized motor-driven bidirectional protein transportation machinery in cilia and flagella.

**Immunoblotting:** Antibody probing protein detection technique involving electrophoretic separation, membrane transfer and detection using target-specific antibody.

**Co-Immunoprecipitation:** Detects protein-protein interaction by pulling down bait proteins with associated targets.

## References

[1] O. P. Ernst, D.T. Lodowski, M. Elstner, P. Hegemann, L.S. Brown, and H. Kandori, Microbial and animal rhodopsins: Structures, functions, and molecular mechanisms, Chem. Rev.114(1), (2014), 126–163. doi: 10.1021/cr4003769.

[2] E. G. Govorunova, O.A. Sineshchekov, H. Li, and J.L. Spudich, Microbial Rhodopsins: Diversity, Mechanisms, and Optogenetic Applications, Annu. Rev. Biochem. 86(1), (2017), 845–872. doi: 10.1146/annurev-biochem-101910-144233.

[3] K. Kojima, A. Shibukawa, and, Y. Sudo, The Unlimited Potential of Microbial Rhodopsins as Optical Tools, Biochem. 59(3), (2020), 218–229. doi: 10.1021/acs.biochem.9b00768.

[4] R. Ochoa-Fernandez, N.B. Abel, F.G. Wieland, J. Schlegel, L.A. Koch, J.B. Miller, R. Engesser et al, Optogenetic control of gene expression in plants in the presence of ambient white light’, Nat. Methods. 17(7), (2020), 717–725. doi: 10.1038/s41592-020-0868-y.

[5] A Reyer, M Häßler, S Scherzer, S Huang, JT Pedersen, KAS Al-Rascheid, E Bamberg et al, Channelrhodopsin-mediated optogenetics highlights a central role of depolarization-dependent plant proton pumps, PNAS. 117(34), (2020), 20920–20925. doi: 10.1073/pnas.2005626117.

[6] Y. Zhou, M. Ding, X. Duan, et al, Extending the Anion Channelrhodopsin-Based Toolbox for Plant Optogenetics, Membranes, 11(4), (2021) 287. doi: 10.3390/membranes11040287.

[7] Y. Zhou, M. Ding, S. Gao, et al, Optogenetic control of plant growth by a microbial rhodopsin, Nat. Plants. 7(2), (2021), 144–151. doi: 10.1038/s41477-021-00853-

[8] K. A. Zalocusky, L. E. Fenno, and K. Deisseroth, 2 Current challenges in optogenetics, in *Optogenetics*. DE GRUYTER, (2013). 23–34. doi: 10.1515/9783110270723.23.

[9] K. Sushmita, S. Sharma, M.S. Kaushik, S. Kateriya, Algal rhodopsins encoding diverse signal sequence holds potential for expansion of organelle optogenetics, Biophys. Physicobiol. 20 (Suppl.) (2023) e201008. 10.2142/biophysico.bppb-v20.s008

[10] R. Tashiro, K. Sushmita, S. Hososhima, S. Sharma, S. Kateriya, H. Kandori, S.P. Tsunoda, Specific residues in the cytoplasmic domain modulate photocurrent kinetics of channelrhodopsin from Klebsormidium nitens, Commun. Biol. 4 (2021) 235. 10.1038/s42003-021-01755-5

[11] J. Wang, D. Deretic, Molecular complexes that direct rhodopsin transport to primary cilia, Prog. Retin. Eye Res. 38 (2014) 1–19. 10.1016/j.preteyeres.2013.08.004

[12] K. Schmidt, et al, Early stages of retinal development depend on Sec13 function, Biol. Open 2 (2013) 256–266. 10.1242/bio.20133251

[13] R. Bhowmick, et al, Photoreceptor IFT complexes containing chaperones, guanylyl cyclase 1 and rhodopsin, Traffic 10 (2009) 648–663. 10.1111/j.1600-0854.2009.00896.x

[14] H. Khanna, Photoreceptor sensory cilium: Traversing the ciliary gate, Cells 4 (2015) 674–686. 10.3390/cells4040674

[15] J.L. Rosenbaum, G.B. Witman, Intraflagellar transport, Nat. Rev. Mol. Cell Biol. 3 (2002) 813–825. 10.1038/nrm952

[16] D.G. Cole, The intraflagellar transport machinery of Chlamydomonas reinhardtii, Traffic 4 (2003) 435–442.

[17] S. Webb, A.G. Mukhopadhyay, A.J. Roberts, Intraflagellar transport trains and motors: Insights from structure, Semin. Cell Dev. Biol. 107 (2020) 82–90. 10.1016/j.semcdb.2020.05.021

[18] E.A. Richey, H. Qin.], Dissecting the sequential assembly and localization of intraflagellar transport particle complex B in Chlamydomonas, PLoS ONE 7 (2012) e43118. 10.1371/journal.pone.0043118

[19] M. Taschner, et al, Crystal structures of IFT70/52 and IFT52/46 provide insight into intraflagellar transport B core complex assembl,. J. Cell Biol. 207 (2014) 269–282. 10.1083/jcb.201408002

[20] L.B. Pedersen, et al, Chlamydomonas IFT172 is encoded by FLA11, interacts with CrEB1, and regulates IFT at the flagellar tip, Curr. Biol. 15 (2005) 262–266. 10.1016/j.cub.2005.01.037

[21] J.A. Follit, et al, The intraflagellar transport protein IFT20 is associated with the Golgi complex and is required for cilia assembly, Mol. Biol. Cell 17 (2006) 3781–3792. 10.1091/mbc.E06

[22] B.T. Keady, Y.Z. Le, G.J. Pazour, IFT20 is required for opsin trafficking and photoreceptor outer segment development, Mol. Biol. Cell 22 (2011) 921–930. 10.1091/mbc.e10-09-0792

[23] M. Awasthi, et al, Novel modular rhodopsins from green algae hold great potential for cellular optogenetic modulation across the biological model systems, Life 10 (2020) 259. 10.3390/life10110259

[24] M. Luck, et a,. A photochromic histidine kinase rhodopsin (HKR1) that is bimodally switched by ultraviolet and blue light, J. Biol. Chem. 287 (2012) 40083–40090. 10.1074/jbc.M112.401604

[25] M. Luck, et a, Photochemical chromophore isomerization in histidine kinase rhodopsin, FEBS Lett. 589 (2015) 1067–1071. 10.1016/j.febslet.2015.03.024

[26] M. Luck, P. Hegemann, The two parallel photocycles of the Chlamydomonas sensory photoreceptor histidine kinase rhodopsin 1, J. Plant Physiol. 217 (2017) 77–84. 10.1016/j.jplph.2017.07.018

[27] Y. Tian, et al, Two-component cyclase opsins of green algae are ATP-dependent and light-inhibited guanylyl cyclases, BMC Biol. 16 (2018) 144. 10.1186/s12915-018-0608-9

[28] T. Pröschold, E.H. Harris, A.W. Coleman, Portrait of a species: *Chlamydomonas reinhardtii*, Genetics (2005) 170:1601–1610.

[29] G.M. Adams, B. Huang, D.J. Luck, Temperature-sensitive, assembly-defective flagella mutants of *Chlamydomonas reinhardtii*, Genetics (1982) 100:579–586.

[30] M.S. Miller, J.M. Esparza, A.M. Lippa, F.G. Lux III, D.G. Cole, S.K. Dutcher, Mutant kinesin-2 motor subunits increase chromosome loss, Mol Biol Cell (2005) 16:3810–3820.

[31] B. Huang, M.R. Rifkin, D.J. Luck, Temperature-sensitive mutations affecting flagellar assembly and function in *Chlamydomonas reinhardtii*, J Cell Biol (1977) 72:67–85.

[32] B.D. Engel, H. Ishikawa, K.A. Wemmer, S. Geimer, K. Wakabayashi, M. Hirono, B. Craige, G.J. Pazour, G.B. Witman, R. Kamiya, W.F. Marshall, The role of retrograde intraflagellar transport in flagellar assembly, maintenance, and function, J Cell Biol (2012) 199:151–167.

[33] W.J. Brazelton, C.D. Amundsen, C.D. Silflow, P.A. Lefebvre, The bld1 mutation identifies the *Chlamydomonas* osm-6 homolog as a gene required for flagellar assembly, Curr Biol (2001) 11:1591–1594.

[34] G.J. Pazour, B.L. Dickert, Y. Vucica, E.S. Seeley, J.L. Rosenbaum, G.B. Witman, D.G. Cole, *Chlamydomonas* IFT88 and its mouse homologue, polycystic kidney disease gene tg737, are required for assembly of cilia and flagella, J Cell Biol (2000) 151:709–718.

[35] K.F. Lechtreck, E.C. Johnson, T. Sakai, D. Cochran, B.A. Ballif, J. Rush, G.J. Pazour, M. Ikebe, G.B. Witman, The *Chlamydomonas reinhardtii* BBSome is an IFT cargo required for export of specific signaling proteins from flagella, J Cell Biol (2009) 187:1117–1132.

[36] S. Sharma, K. Sushmita, R. Singh, S.K. Sanyal, S. Kateriya, Phototropin localization and interactions regulate photophysiological processes in Chlamydomonas reinhardtii, Biochimie (2025). 10.1016/j.biochi.2025.08.014

[37] M. Awasthi, et al, The trafficking of bacterial type rhodopsins into the Chlamydomonas eyespot and flagella is IFT mediated, Sci. Rep. 6 (2016) 34646. 10.1038/srep34646

[38] D.G. Cole, D.R. Diener, A.L. Himelblau, P.L. Beech, J.C. Fuster, J.L. Rosenbaum, Chlamydomonas kinesin-II-dependent intraflagellar transport (IFT): IFT particles contain proteins required for ciliary assembly in Caenorhabditis elegans sensory neurons, J Cell Biol (1998) 141:993–1008

[39] E. Betleja, D.G. Cole, Ciliary trafficking: CEP290 guards a gated community, Curr. Biol. 20 (2010) R928–R931.

[40] B. Craige, C.-C. Tsao, D.R. Diener, Y. Hou, K.-F. Lechtreck, J.L. Rosenbaum, G. B. Witman, CEP290 tethers flagellar transition zone microtubules to the membrane and regulates flagellar protein content, JCB (J. Cell Biol.) 190 (2010) 927–940.

[41] S.A. Baker, et al, IFT20 links kinesin II with a mammalian intraflagellar transport complex that is conserved in motile flagella and sensory cilia, J. Biol. Chem. 278 (2003) 34211–34218. 10.1074/jbc.M300156200

[42] A. Chien, et al, Dynamics of the IFT machinery at the ciliary tip, eLife 6 (2017) e28606. 10.7554/eLife.28606

[43] D. Szklarczyk, et al, STRING v11: protein–protein association networks with increased coverage, supporting functional discovery in genome-wide experimental datasets, Nucleic Acids Res. 47 (2019) D607–D613. 10.1093/nar/gky1131

[44] Fu H, Subramanian RR, Masters SC (2000) 14-3-3 proteins: structure, function, and regulation. Annu Rev Pharmacol Toxicol 40:617–647

[45] Yaffe MB, Rittinger K, Volinia S, Caron PR, Aitken A, Leffers H, Gamblin SJ, Smerdon SJ, Cantley LC (1997) The structural basis for 14-3-3: phosphopeptide binding specificity. Cell 91:961–971

[46] Johnson C, Crowther S, Stafford MJ, Campbell DG, Toth R, MacKintosh C (2010) Bioinformatic and experimental survey of 14-3-3-binding sites. Biochem J 427:69–78

[47] Coblitz B, Shikano S, Wu M, Gabelli SB, Cockrell LM, Spieker M, Hanyu Y, Fu H, Amzel LM, Li M (2005) C-terminal recognition by 14-3-3 proteins for surface expression of membrane receptors. J Biol Chem 280:36263–36272

[48] Ganguly S, Weller JL, Ho A, Chemineau P, Malpaux B, Klein DC (2005) Melatonin synthesis: 14-3-3-dependent activation and inhibition of arylalkylamine N-acetyltransferase mediated by phosphoserine-205. Proc Natl Acad Sci USA 102:1222–1227

[49] S. Sharma, K. Sushmita, S.K. Sanyal, I. Sizova, P. Hegemann, S. Kateriya, Molecular characterization of phototropin signalosome regulating photophysiological processes in green algae, Biophys. J. 122 (2023) 466a. 10.1016/j.bpj.2022.11.2499

[50] S. Sharma, S.K. Sanyal, K. Sushmita, M. Chauhan, A. Sharma, G. Anirudhan, et al, Modulation of phototropin signalosome with artificial illumination holds great potential in the development of climate-smart crops, Curr. Genomics 22 (2021) 181–213. 10.2174/1389202922666210412104817

[51] S.K. Sanyal, M. Awasthi, P. Ranjan, S. Sharma, G.K. Pandey, S. Kateriya, Characterization of Chlamydomonas voltage-gated calcium channel and its interaction with photoreceptor support VGCC modulated photobehavioral response in the green alga, Int. J. Biol. Macromol. 245 (2023) 125492. 10.1016/j.ijbiomac.2023.125492

[52] K. Sharma, I. Sizova, S.K. Sanyal, G.K. Pandey, P. Hegemann, S. Kateriya, Deciphering the role of cytoplasmic domain of channelrhodopsin in modulation of the interactome and SUMOylome of Chlamydomonas reinhardtii, Int. J. Biol. Macromol. 243 (2023) 125135. 10.1016/j.ijbiomac.2023.125135

[53] S.K. Sanyal, K. Sharma, D. Bisht, S. Sharma, K. Sushmita, S. Kateriya, G.K. Pandey, Role of calcium sensor protein module CBL-CIPK in abiotic stress and light signaling responses in green algae, Int. J. Biol. Macromol. 237 (2023) 124163. 10.1016/j.ijbiomac.2023.124163

