## Supplementary materials for "The protein turnover and trafficking of Chlamyopsin6 is regulated by IFT88 and IFT52 in the *Chlamydomonas reinhardtii*"

**Short running title:** Intraflagellar transport-mediated trafficking of modular rhodopsins in *Chlamydomonas reinhardtii*

Kumari Sushmita^1#^, Sunita Sharma^1,2#^, Rajani Singh^1^, Suneel Kateriya^1*^

^1^Laboratory of Optobiotechnology, School of Biotechnology, Jawaharlal Nehru University, New Delhi 110067, India;

^2^Department of Cellular Biology, University of Georgia, Athens 30602, Georgia, USA.

^#^The authors contributed equally to this work.

Orcid ID: Kumari Sushmita (**0000-0001-9649-1446), Sunita Sharma (0000-0002-6531-7522), Rajani Singh (**0000-0002-1520-2794), **Suneel Kateriya (0000-0001-5428-4297)**

**Supplementary Tables**

**Table S1: Brief technical description of algal strains used in this study.**

| Strains (Wild  type/Mutants) | Generation Method and brief description | Growth Temperature | Reference |
| --- | --- | --- | --- |
| CC-124 and CC- 125 | This is the basic “137c” wild type strain. CC-  124 and CC-125 carry the nit1 and nit2 mutations, and cannot grow on nitrate as their sole N source. CC-125 carries the AGG1 (agg1+) allele for phototactic aggregation, in contrast to CC-124, which has the agg1 allele at this locus. | 22˚C | [28] |
| CC-1396 fla8 mt | This is a conditional mutant (temperature- sensitive). The fla8 mutant fails to  assemble flagella at 34 degrees.  In CC-1396, the glutamic acid at position 21 is changed to a lysine (E21 to K21 ) via a G-to-A transition in FLA8 locus. | Permissive 22 ˚C  Non-permissive 33 ˚C | [29]  [30] |
| CC-1919 fla10-1 mt- | The fla10 mutant cannot regenerate flagella at 32 degrees. The original fla10 allele (fla10-1) is a C to A transversion that alters amino acid 329 in the motor domain of the kinesin-homologous KHP1 protein. | Permissive 22 ˚C  Non-permissive 33 ˚C | [30,31] |
| CC-4424 dhc1b- 3 fla10 | This strain was generated by mating dhc1b-3 mt+ (CC-4422) to fla10 mt-. It is temperature sensitive, losing its flagella at 34ûC with similar kientics to fla10.  CC-4422 is a strain having an A to T transversion at position 12313 of the DHC1b coding sequence that results in an Ile to Phe substitution at amino acid 4105. | Permissive 22 ˚C  Non-permissive 33 ˚C | [32] |
| CC-477 bld1-1 | This is the original bld1 mutant shown to have deletion in gene encoding IFT52  The BLD1 locus encodes intraflagellar transport protein IFT52, a homolog of C. elegans osm-6. | 22˚C | [33] |
| CC-3943 ift88- 1::NIT1 mt+ | This is Pazour strain V79, the original ift88-1  mutant isolate. This is generated by random insertional mutagenesis. | 22˚C | [34] |
| CC-1920 fla11  mt- | The fla11 mutant slowly resorbs flagella at 32 degrees and is slow to generate flagella when induced to form gametes at 21 degrees. This is a point mutation in the gene encoding the IFT172 protein, resulting in a Leu to Pro change at amino acid residue 1615. | Permissive 22 ˚C  Non-permissive 33 ˚C | [20] |
| CC-4374 cep290-1 | This is a mutant in which most of the CEP290 gene has been deleted.  Mutant cells are mostly palmelloid and have very short/stumpy flagella once released from the mother cell wall by treatment with an autolysin. | 22˚C | [35] |

**Table S2: Accession number and sequence of modular rhodopsin used in this study**

| **Name** | **Accession number** | **Sequence** |
| --- | --- | --- |
| **Cop5** | AAQ16277.3 | MAPTGSLPSHLQDRIDALETNQRDTDAQAFQKAVERRARVSFSVAIAGFGVLLYLLASYLICTSNLGDPELTAQFHAQADPNAYTWPMVAFGTAFGLNFITLLFERESAKFQLALLACYINFLAGFSDYMSWKGYAPIVRDSWGQGFQLLRTVMWLLTTPAMVYLLSIISDFSRLKVYSVMLADVLMITFGILAFLAYNKVMSILFYMVAWCLFAYVVHSMWSMFHASIAEARHDSSRVSLEVLRLFAVGLWFTFPVIWIVVKMGLVDIRTEEWTWCACDFLGKVMFSSSLLHGNFLTIEQRRLIAMRIVEEGNRIQVIQELKDLVEQKERFMSSMSHELRTPLNGIIGLSDALLVGSCGEINDQALKTITTIKTSGARLLNLINDILDAASMRKGKLTIKHEKVNLKRVVDDVIDLCQPLAKRGVKLVNDLRENVPFVLGDTGRIIQVFHNLIGNSCKFTHSGNISISAAVKDDEVEVAVADTGIGIPEDKFDQIFLAFEQVDMSVTRKYGGTGLGLNLVKQLVEAHGGRISVKSRENQGTTFYFTLKIHSEHPNEGQPQMTPTESVAAIPPPEHSQLATVRGTAPSAGGLPPSSSGATAGPRRAPSRRGSFTDKMLGALGAGMGGHHKKPPSETGKSSLGPHGGAGAGAGGAGAGTAAAAAGGSAGYASAAAPRPSQTPSSPSAAVATAAAAGQHDDRESRDGPYASNAGPGGGGNAGQGSAMVRRVSRDAHGSNPPNPKSVTAGPQVSGPMSLSDAAALMKGALKRKSSFRERGAKVRVLSVDDDPVNQLVIQNLLAPVGYEILQAMDGQEALKVLTEEERLPDVILLDVMMPGMSGYEVCRKLREMYPLSCIPVIMISAKSKEEHIVEGLAAGSNDYVVKPFGRQEILARIAAHLRFRDTVYQAGEIAGAIPGDVRPERVLLRGGANGGGLDLTPFLTGPARFTSLPPRIARGIEAGTTSTTLQMFESLTLLEVRLVNLGDLLASVPASDLLVALASLFHDLDTLLEQHGCYLLEGLDESHLIVSGLDNVGDQVLHALGLARSLIAAADTFALGGRRSKLHLAVGVHTGPAQGVLVGYSHPLIFFTGQLPAEVHMLQATCPPNCVHVSARVLESVAHSEREHFVPAGVMASGATTYLMKVGGWEGGGIVAASEATSRWSKAKRGMHDGLATDARRMRPIQLALLDMAQANPAFLVDIASAAHGAAGGGAGTSSDGQGERGGGGGGSEELSKLKREYESLERQMEEVSGEAARLQDLVDELEEQLLARSNTAAASTAATAAASQQAAVAAADAASLKAHVAELEGQLAEAARERNGLEAALAEMEQRLVATHAALATANHEAADRARAHTPAPQELPPHHPHSASRSHLHVLPQQNTPRVETASMAPSASGSEANFASGLSALGGGSLFGGGALMPRGAMPLGVPMRMPNLAALRAHAGDMRGLLDELGLGSLASRFEAEDVSPGLLPYLDDAAMRELGASSVGARLKLRLAAQALFM* |
| **Cop6** | Cre11.g467678.t1.1 | MKLRQRTVGAQLRSQPVSSAGGPANSGPPATPSGGIAPVSIFGAAEALADPEARGWILTWSWTFTGFFVYITASWLSGLWYTTDPLAYAALRAQVPTLVYQMSSTAFFTALVLNLTSLLFEDNAPKRQLALLSCAIKGAACHTDLLLVTGGATVLYDAYGSICIPQRYVQWLVTTPTMVYILSKISDFTPRQTATAIGLDVLMVLSGLVANFLRSPYLWVAFLTSTAAFIGVLYMMGLMVYSAVKEHTSANSRRSLLFIYMCTLFIWNLFPLAWILHVVHRGSPAAEYLNVFANFMAKVLFSSSIMYGNYMTIAQRRLLAQQDAENANRVQMIQDLRDSVTRKDQFMSLMSHELRTPLNGIIQLSDALVRGAGGEMNPKGQHFVRTIKNSSNHLLNIINDILDVAALKEGKLTIKHEVCSLAKAVDHVVDIVAPLAKKEVTMERWVDPATPLIIADFSRVIQILYNLTGNALKFTNKGRVGVRVEPSADGTHVLLQVSDTGIGIPKDRLHSIWGAFEQVDMSVTRKYGGTGLGLNIVKQLVEAHEGTIEVASVEGRGTTFTVELPVLQSSTRRSLEGQVLDSLTRCGHAAARDTMVQRRTRSRPSLGLEDTITQFARGVQRRASGLLGVAKAAQEAGSNAGTGPAGSTGAGAGGGGGAGGGGQRDSLDREEGEQLLRKRTHELEHESRLGDLARRTVHKQSMEEADLASRQLLLSDYERRAERDTRSLERIDSGQPGLTNGGGEGAGGAAGGGAGSGGGGGSAPGSAGKAGVAGGSCRGDVRGGGTDGRGGGGGGAGGAGGGGGGGNGGGGASGGGGRRSGAPTSGRASLGELPSGGSGGGGGGGGGGGDGTPESPRSAIARRGLLAMRQSSLSNLRGATSARAGGAAGGAAASGGGVSVGRWASTTDTGFANQGAAAAAWRGDSHRTLPGVEGGCVSVSSANGNSIADVYEALQLARASNESGGGGGGGGGGGGGGNGSSLKASGGSLALRMSAYGRTGYGGANGGGGGGANGLNGYGGGGGALGGSGASGSLLKSALESDLYRNGPSPRDPYDCDASSVGADSEYEWVGDGGGAGGGGAFGGRSRGPTSTGSLALGGMACGPGGRRHQPPKPSLRSGKLTPMLSNAAAAAAAITAVPPGLTLDKLAYSDMYGTIQVLSVDDEDINQIVLEEILTDSGYAFARCMDGAEALEWLCASDTMPDLILLDCMMPVMSGHEFCATLRKVIPGNVLPVIMVSAKSDEENIVEGLRSGSNDFVRKPYQREELLARIETQLRLKSDSWWLAELVNNVDGRETESMKLLKNILPESIIARMQQGQKFVADSHGHVVILFSDIVGFTSLSSKLPTAEVFLMLSNMFTAFDKLTDRFSVYKVETIGDAYMVAAGHDEDEDKEAKGSPLMRVLGFARAMLDVVRNITAPNGERLRIRIGVHCGPAFAGVIGMKCPRYCFLGDTVNTASRMESTGFPMCIHVSENVFKHHPAAEAELQEVGERDIKGKGHMRTYVVRTGAWEQALRDFAARQQAAAAAAQAQAQTLALARQQAALLQQQHEQLQLQLQANGGGGGGGATANAAKAAADALAQLPASLSFDSASAGSTSSMLPLSGGSVAGAAARTANSLAPGAGGGGAAAGAADGGGGGNGCGGGGGAVRMGHLLSTVAEEGSLPGSPSSFAAAAAAAALSPSAAAQQQQQQAPGYHPSHLRTASAGAGRSPLSRGELPAAASSPLQPGGGAAAGAAGSSPLLPPLLRGANHYTDPRLAGGLRPSFSTGGYVLEDDSDDGGTSTNTGVGGGTGHGMGSSLGEGRAGSISLTGIGARSFTGASCGGGAGGGGGGGGLRMLNSPTGQYDTGDAGENGSSGPGGSGGGGGEVGGSGGTPSGLRVHVPRSLSGLSGGSYGSGGSTGIGMSRAGLGSATAAAAAASGACFLTSDAADASFSGGVLAATTSHAALGSGGAADSQHAPSAAGAVMGSSPAAALQVSPASPGGAAAGGLGSGLSPQSSLYGGAQLLLSPNSAGLHSGHANTTLAYLEQRIASLGTQLATEALSRQRLQDELDAERRRAAGAMQQASLLMQQLRNATQAAGGGSGGAPSSVAAAAAAAGQEQHLARLQLGSGGGGADAASGGGGGAALPPSAAVPLSRLPPPPRGAGGAVASLRAAGGANELLPTSNEANSNADADVIIGGGGMSSTVVPAAQPPSSSAFGSAGAATSGGSGNDDQEITLASPFVSDLPAYALEAGDVVPNSLDFASQGFDPGSSGAGGRGGGGGSRHAGRAAAAQQRPSSKVQRMNVGDAAALMLQPPKAQQLQPGSQSTAFVEAGGVGRSGASGGGIAEPLPGSAAEAYSGYSRQQLEDQLLLPVPMPTSITALMPGGGTAAATAAVGTATAASTTTTTTTTVSHHRHVFHTSAAAGLALSAHPAPGSSPLPAPSSSSPCTALQPLQLQPPPLPCYSLDALFVDLGLEPYLPRFRDEAIRLDMLLSMDAQQLERLGLKPLGYRIRVREAVVELARGLLRSSEDAAVLVEMSQQHQHQQQYRSQQAQQLQQQHQPLER* |

**Table S3: List of different nodes with their description involved in the predicted protein-protein networking**

| **Node** | **Description** |
| --- | --- |
| FAP116 | *Flagellar associated protein. (510 aa)* |
| CAM1 | *Calmodulin; Belongs to the calmodulin family. (163 aa)* |
| STM2 | *Ubiquitin-like domain-containing protein. (153 aa)* |
| PKL1 | *Serine/threonine-protein phosphatase. (890 aa)* |
| ARL2 | *Uncharacterized protein; Belongs to the small GTPase superfamily. Arf family. (168 aa)* |
| A0A2K3CVV1 | *Small ubiquitin-related modifier. (106 aa)* |
| IFT80 | *Intraflagellar transport protein 80. (765 aa)* |
| A0A2K3D7N7 | *Chlamyopsin 5 (2549 aa)* |
| PHOT | *Phototropin; Protein kinase that acts as a blue light photoreceptor. Required for non-photochemical quenching (NPQ), a mechanism that converts and dissipates the harmful excess absorbed light energy into heat and protect the photosynthetic apparatus from photo-oxidative damage. Controls the energy-dependent chlorophyll fluorescence quenching (qE) activity of chlorophyll excited states by inducing the expression of the qE effector protein LHCSR3 in high light intensities. (749 aa)* |
| A0A2K3DIH9 | *Uncharacterized protein. (3106 aa)* |
| FLA8 | *Kinesin-like protein; Belongs to the TRAFAC class myosin-kinesin ATPase superfamily. Kinesin family. (725 aa)* |
| DHC1b | *Uncharacterized protein. (4333 aa)* |
| BLD1 | *Intraflagellar transport protein IFT52. (454 aa)* |
| A0A2K3DMV5 | *Uncharacterized protein. (2729 aa)* |
| CPH1 | *Cryptochrome photoreceptor. (1008 aa)* |
| EB1 | *Microtubule plus-end binding protein. (280 aa)* |
| PPA1 | *Serine/threonine-protein phosphatase. (307 aa)* |
| UBC3 | *E2 ubiquitin-conjugating enzyme; Belongs to the ubiquitin-conjugating enzyme family. (163 aa)* |
| UVR3 | *Cryptochrome photoreceptor. (595 aa)* |
| CPKG1 | *cGMP-dependent protein kinase. (1027 aa)* |
| cop4 | *Chlamyopsin 4 light-gated ion channel. (737 aa)* |
| IFT172 | *WD_REPEATS_REGION domain-containing protein. (1755 aa)* |
| A0A2K3DFH6 | *ThiF domain-containing protein. (391 aa)* |
| IFT81 | *IFT81_CH domain-containing protein. (683 aa)* |
| A0A2K3DQP2 | *Uncharacterized protein. (750 aa)* |
| IFT88 | *Intraflagellar transport particle protein 88. (782 aa)* |
| COP3 | *Uncharacterized protein. (719 aa)* |
| MKP1 | *Uncharacterized protein. (553 aa)* |
| A8J3A7_CHLRE | *Small ubiquitin-related modifier. (97 aa)* |
| FAP259 | *Flagellar associated protein. (647 aa)* |
| FLA10 | *Kinesin-like protein; Belongs to the TRAFAC class myosin-kinesin ATPase superfamily. Kinesin family. (786 aa)* |
| DYF13 | *Uncharacterized protein. (554 aa)* |
| NIT1 | *Nitrate reductase; Nitrate reductase is a key enzyme involved in the first step of nitrate assimilation in plants, fungi and bacteria. (882 aa)* |
| A0A2K3CPY7 | *Uncharacterized protein. (562 aa)* |
| A0A2K3E007 | *Chlamyopsin 5 (1501 aa)* |
| FAP22 | *Uncharacterized protein. (443 aa)* |
| FTT1 | *14-3-3 protein; Belongs to the 14-3-3 family. (259 aa)* |
| A0A2K3CPZ6 | *Uncharacterized protein. (648 aa)* |
| IFT20 | *Intraflagellar transport particle protein IFT20. (135 aa)* |
| IFT57 | *Uncharacterized protein. (469 aa)* |
| IDA5 | *Actin. (377 aa)* |
| IFT74 | *Uncharacterized protein. (641 aa)* |
| A8J3A8_CHLRE | *Small ubiquitin-related modifier. (94 aa)* |

**Supplementary Figures:**

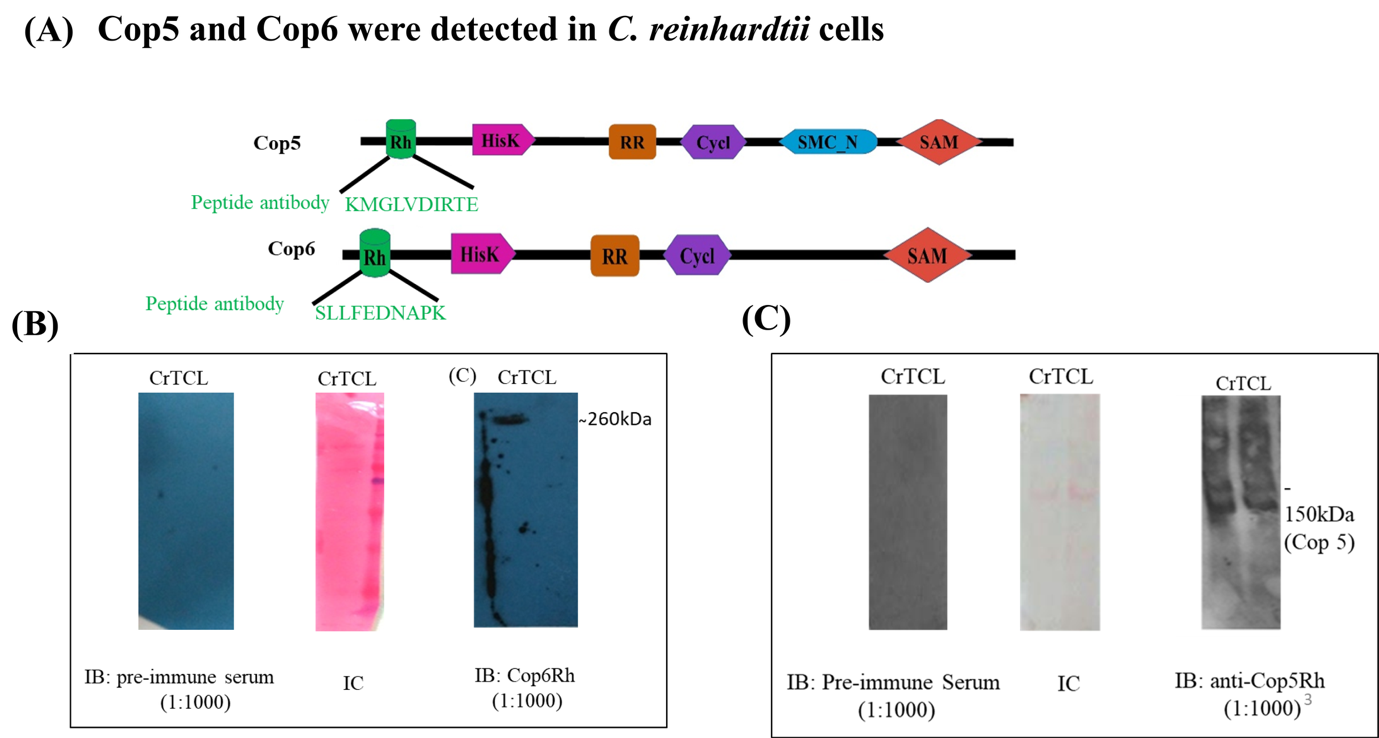

**Figure S1:** **The conserved domain architecture and detection of Chlamyopsin5 (Cop5) and Chlamyopsin6 (Cop6) in the *Chlamydomonas* cell lysate.**

(A) Represents the conserved domain architecture of Cop5 and Cop6; (B) and (C) represents the detection of Cop6 and Cop5 in the cell lysate of *Chlamydomonas* using immunoblotting*.* The molecular weight marked represents the calculated molecular weight of the specific protein. Pre-immune serum as a negative control, IC-Internal Control, where Cop5Rh and Cop6Rh are specific antibodies against the peptides of the rhodopsin domains Cop5 and Cop6,
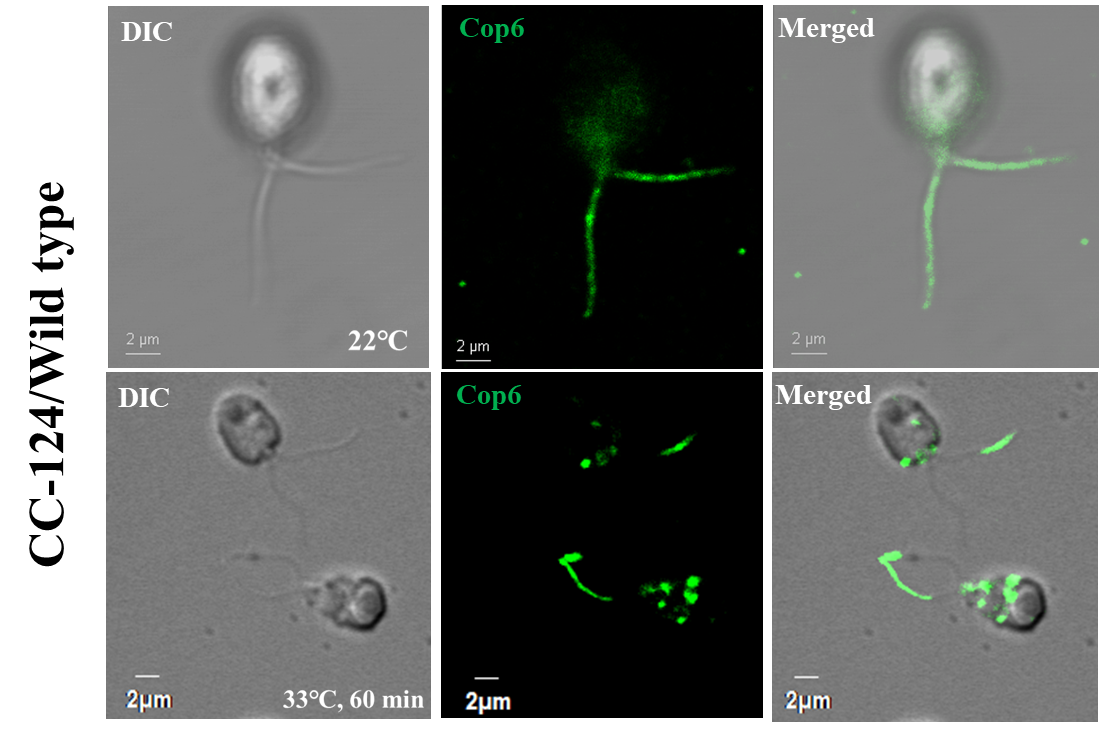
respectively.

**Figure S4: Immunolocalization of Cop6 in wild type of *C. reinhardtii* cells at 22℃ and 33°C temperature.** The first panel represents immunostaining of cells for Cop6 (Green) with anti-Cop6 (1:250 dilution). The secondary antibody for Cop6 is Alexa 488 conjugated with anti-goat IgG (1:1000 dilution). The first panel represents DIC, third panel represents signal for Cop6 and third panel show an overlay of the second panelwith DIC. Scale bar=2 μm.
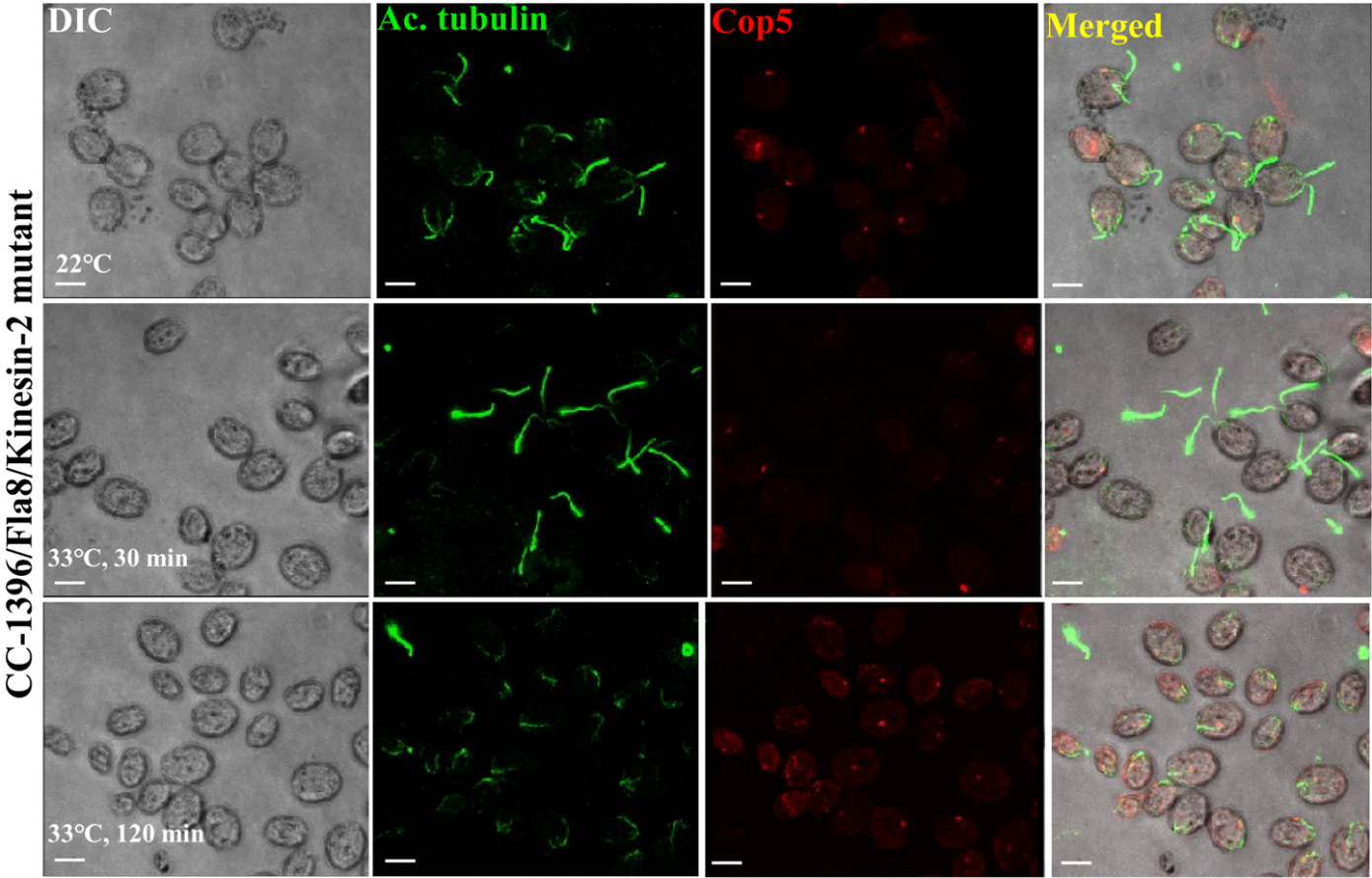

**Figure S2: Immunolocalization of Cop5 in kinesin motor mutant strain (CC-1396) of *C. reinhardtii* cells**.

Cop5 localization to the eyespot is unperturbed in the kinesin motor mutant strain at a non-permissive temperature. The first panel represents DIC, and the second panel represents the signal for acetylated tubulin (1:1000 dilution) (green). Alexa 488 conjugated anti-mouse IgG was used as the secondary antibody. The third panel represents the signal for Cop5 with anti-Cop5 (1:250 dilution) (Red). Alexa 546 conjugated anti-rabbit IgG was used as the secondary antibody. The fourth panel is an overlay of the second and third panels with DIC. Scale bar = 5 μm

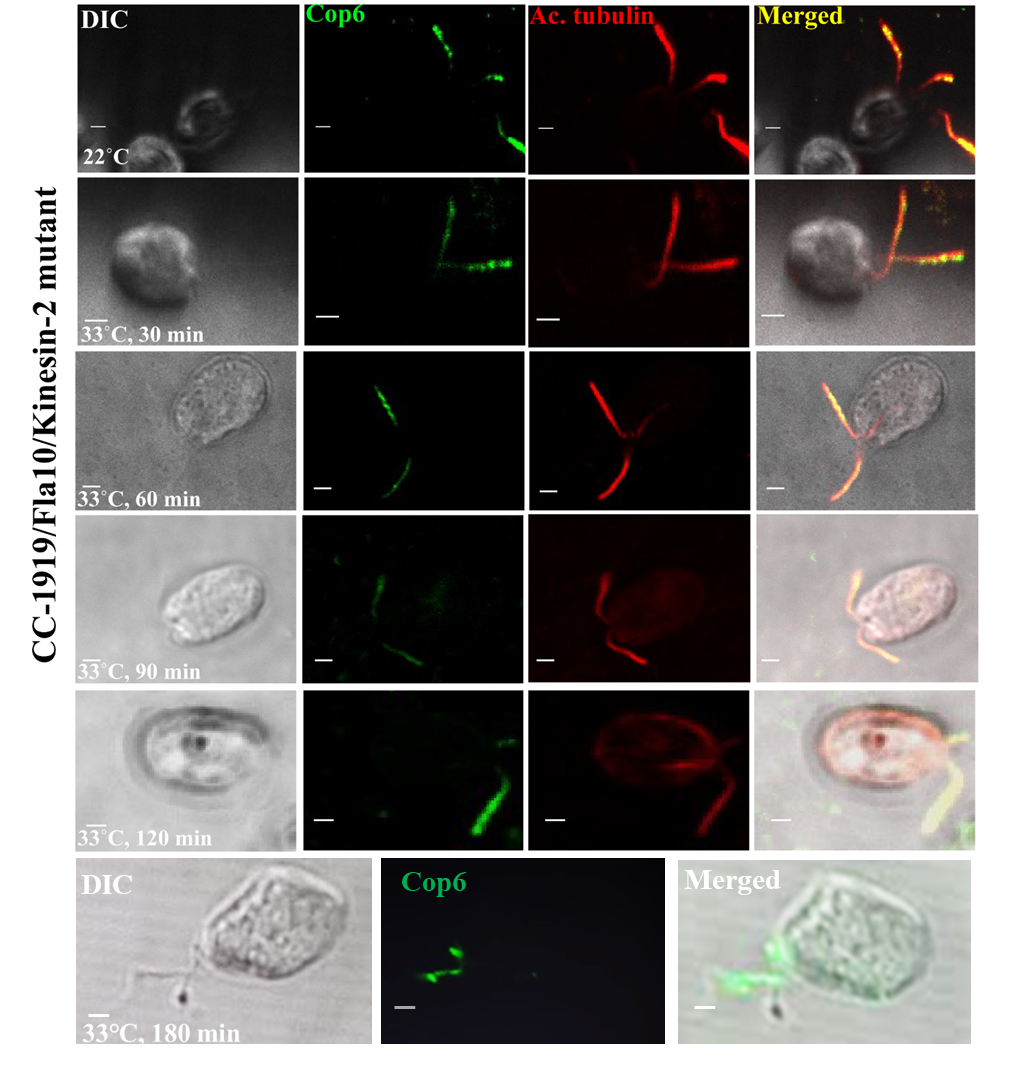

**Figure S3: Immunolocalization of Cop6 in the Kinesin2 mutant strain (CC-1919) of *C. reinhardtii* cells**. Immunostaining of cells was performed using anti-acetylated tubulin (1:1000) (Red) and anti-Cop6 (1:250) (Green). The secondary antibody (1:1000 dilution) for acetylated tubulin and Cop6 was Alexa 647 and Alexa 488 conjugated anti-mouse IgG and anti-goat IgG, respectively. The first channel shows DIC and the second panel depicts the fluorescence specific for Cop6, the third for acetylated tubulin. The fourth panel depicts an overlay of the second and third panels with DIC. Scale bar=2 μm.

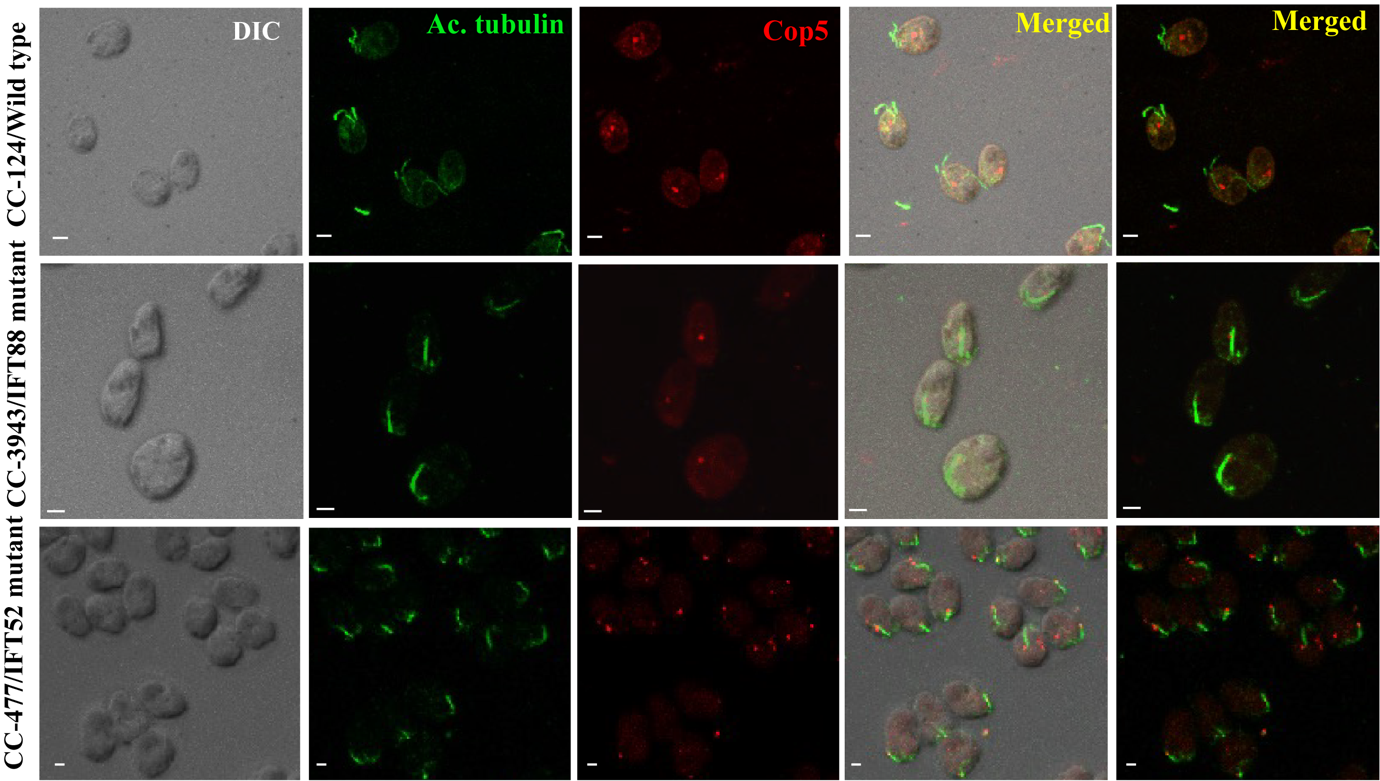

**Figure S5: Immunolocalization of Cop5 in IFT52 and IFT88 mutant strain of *C. reinhardtii* cells**. Cop5 trafficking remains unaffected by IFT disassembly and destabilization of IFT52 and IFT88. Cells were stained with acetylated tubulin, dilution 1:1000, (green, second panel) followed by a second labelling with anti-mouse IgG conjugated with Alexa 488 (1:1000 dilution). Cop5 was labelled with anti-Cop5, dilution 1:250, (red, third panel), followed by a second labelling with anti-rabbit IgG conjugated with Alexa 546 (1:1000 dilution). The first channel shows DIC. The fourth and fifth panels depict an overlay of the second and third panels with and without DIC, respectively. As these are not temperature sensitive mutants, the temperature and time point is same as wild-type i.e. at 22℃. Scale bar=2 μm.

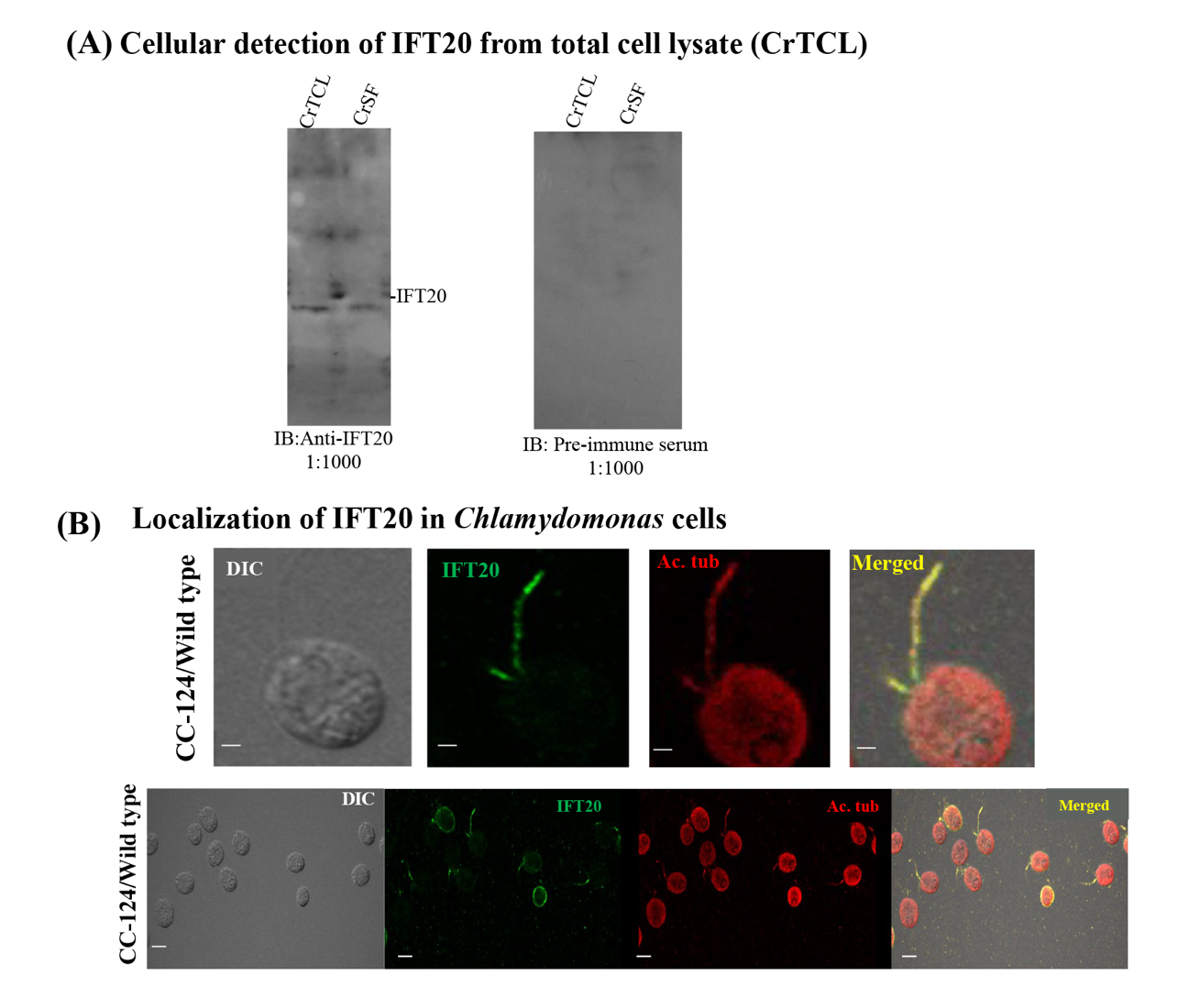

**Figure S6: Immunoblotting and localization of IFT20 in *Chlamydomonas* cells.**

IFT20 was seen to be localized in the flagella of *Chlamydomonas* cells. (A) Represents the detection of IFT20 in the cell lysate of *Chlamydomonas* using immunoblotting*.* Pre-immune serum is a negative control, whereas anti-IFT20 is a specific antibody against the full-length recombinant IFT20. (B) Represents the localization of IFT20 in *Chlamydomonas* cells. The first panel depicts DIC and the second panel depicts a signal for IFT20, the third is for acetylated tubulin, and the fourth panel represents the overlay of the second and third panels with DIC. The upper panel and lower panel show single cells and a group of cells, respectively. Scale bar = 2 μm in a single cell (upper panel) and scale bar = 5 μm in a group of cells (lower panel). CrTCL: *Chlamydomonas reinhardtii* total cell lysate (fraction obtained after cell lysis); CrSF: *Chlamydomonas reinhardtii* soluble fraction (supernatant collected after centrifugation of the cell lysate).

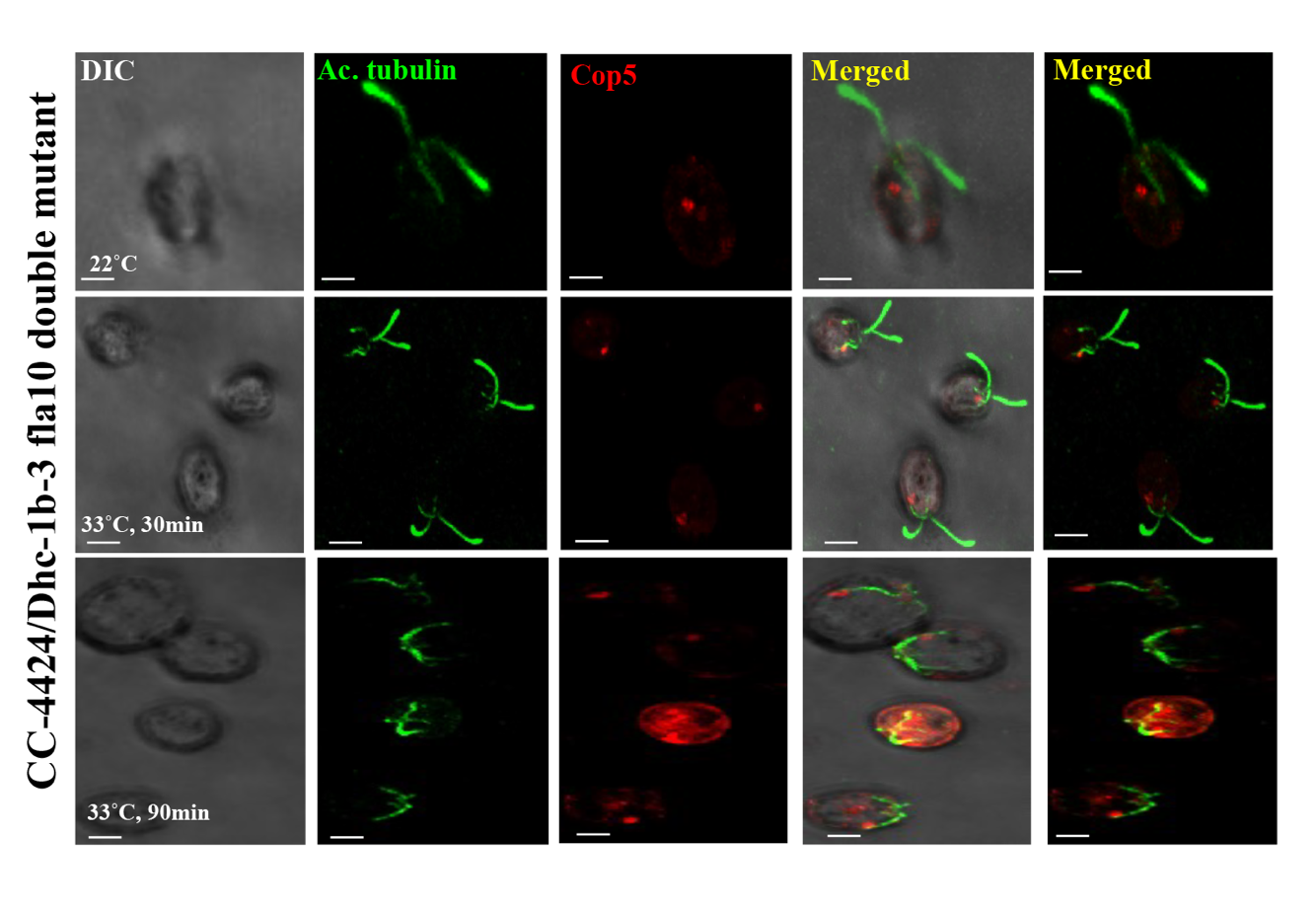

**Figure S7: Immunolocalization of Cop5 in Dynein heavy chain motor (Dhc-1b-3) and kinesin-2 (fla10) double mutant strain (CC-4424) of *C. reinhardtii cells*.**

Anterograde and retrograde movement of Cop5 within the cell body is not controlled by Kinesin and Dynein motors. Flagella of the cell were stained with anti-actylated tubulin (1:1000 dilution) in green and Cop5 (Red) with anti-Cop5(1:250 dilution), secondary antibody for acetylated tubulin and Cop5 is Alexa 488 and Alexa 546 conjugated anti-mouse and anti-rabbit IgG, respectively (1:1000 dilution). The first panel represents DIC, and the second panel shows the signal for acetylated tubulin; the third panel is for Cop5. The fourth and fifth panel is an overlay (merged) of the second and third panels with and without DIC, respectively. Scale bar = 5 μm.

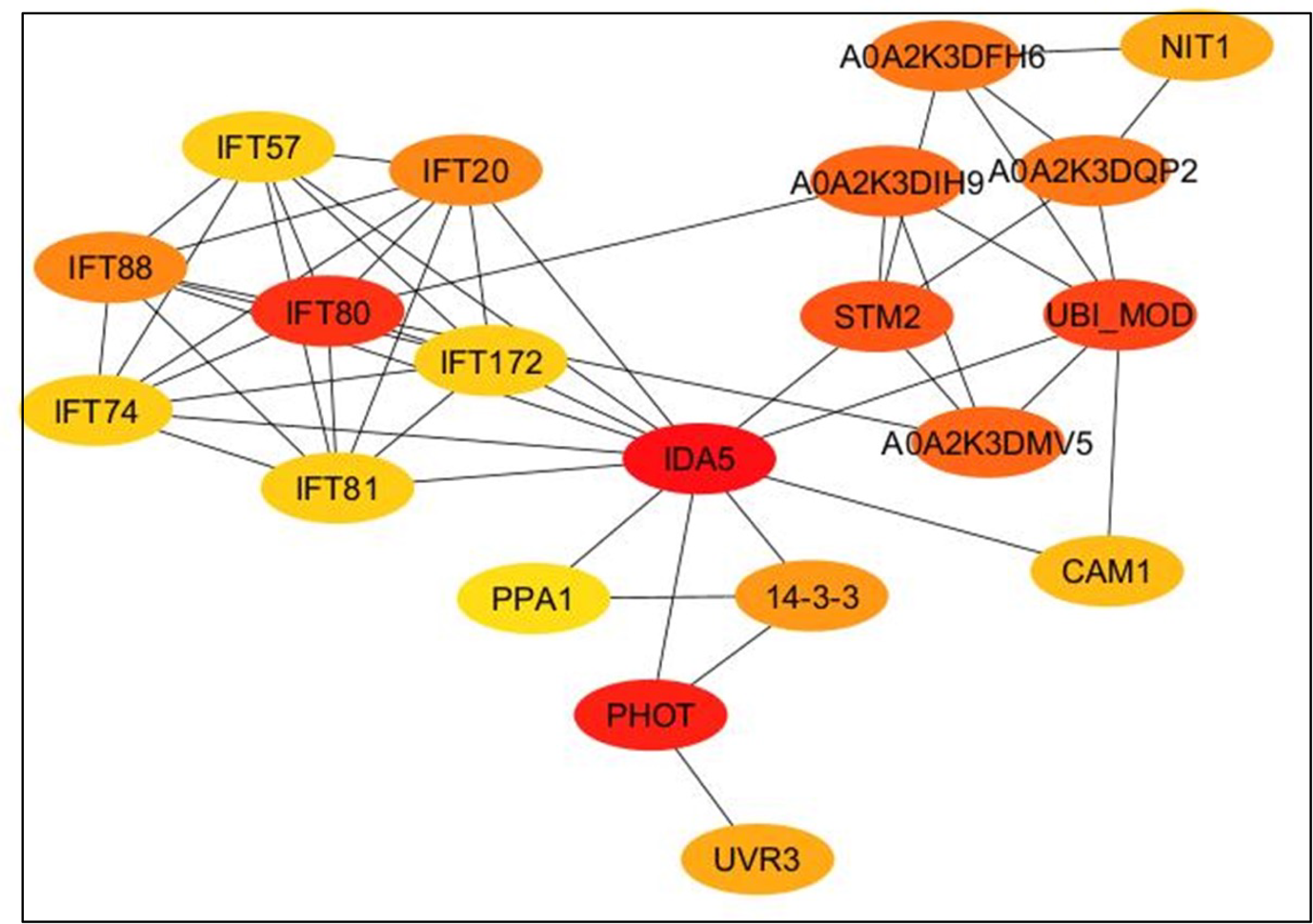

**Figure S8: Represents the important nodes modulating the predicted interactome shown in Figure 8.**

The analysis was performed in Cytoscape to identify hub modules using the cytohubba plugin. The top 20 betweenness algorithms and shortest pathways were used using the default settings. The top 20 rank was based on the betweenness score, shown in colour code from red to orange.
