## Appendix for "The protein turnover and trafficking of Chlamyopsin6 is regulated by IFT88 and IFT52 in the *Chlamydomonas reinhardtii*"

**Short running title:** Intraflagellar transport-mediated trafficking of modular rhodopsins in *Chlamydomonas reinhardtii*

Kumari Sushmita^1#^, Sunita Sharma^1,2#^, Rajani Singh^1^, Suneel Kateriya^1*^

^1^Laboratory of Optobiotechnology, School of Biotechnology, Jawaharlal Nehru University, New Delhi 110067, India;

^2^Department of Cellular Biology, University of Georgia, Athens 30602, Georgia, USA.

^#^The authors contributed equally to this work.

Orcid ID: Kumari Sushmita (**0000-0001-9649-1446), Sunita Sharma (0000-0002-6531-7522), Rajani Singh (**0000-0002-1520-2794), **Suneel Kateriya (0000-0001-5428-4297)**

**Supplementary protocols**

**Protocol S1: Generation of antibodies against Cop5 and Cop6**

The Cop5 and Cop6 antibodies raised against the peptides of ten amino acids of the rhodopsin domain were provided by Dr. Peeyush Ranjan. The generalized protocol followed for raising the antibodies against the peptide is discussed below.

Amino acid sequences of the Rhodopsin domains of Chlamyopsins from *Chlamydomonas reinhardtii* were aligned to select peptides of 10-15 amino acids unique to Cop5 and Cop6 using a web-based tool (http://multalin.toulouse.inra.fr/multalin/). The antigenicity of the selected peptides was predicted via a web-based immunoinformatics tool (<http://www.iedb.org/>). The commercial facility (Merck-Bangalore Genei, India) was utilized to raise primary antibodies against the selected peptides, KMGLVDIRTE for Cop5 and SLLFEDNAPK for Cop6, in rabbit and goat, respectively. The dilution of antibodies used for immunoblotting is 1:1000, and immunostaining is 1:250

**Protocol S2: Generation of an antibody against IFT20.**

Dr. Peeyush Ranjan kindly provided the IFT20 gene cloned in the pET21a vector. The full-length recombinant IFT20 protein was expressed and purified. The purified protein was utilized for raising a primary antibody in a goat through a commercial facility, Genei Services (Sirifort Associates). Prior to the finalization of the host, the cross-reactivity was analyzed via immunoblotting of Chlamydomonas total cell lysate (CrTCL) with the host serum. The dilution of antibodies used for immunoblotting is 1:1000 and immunostaining is 1:250.

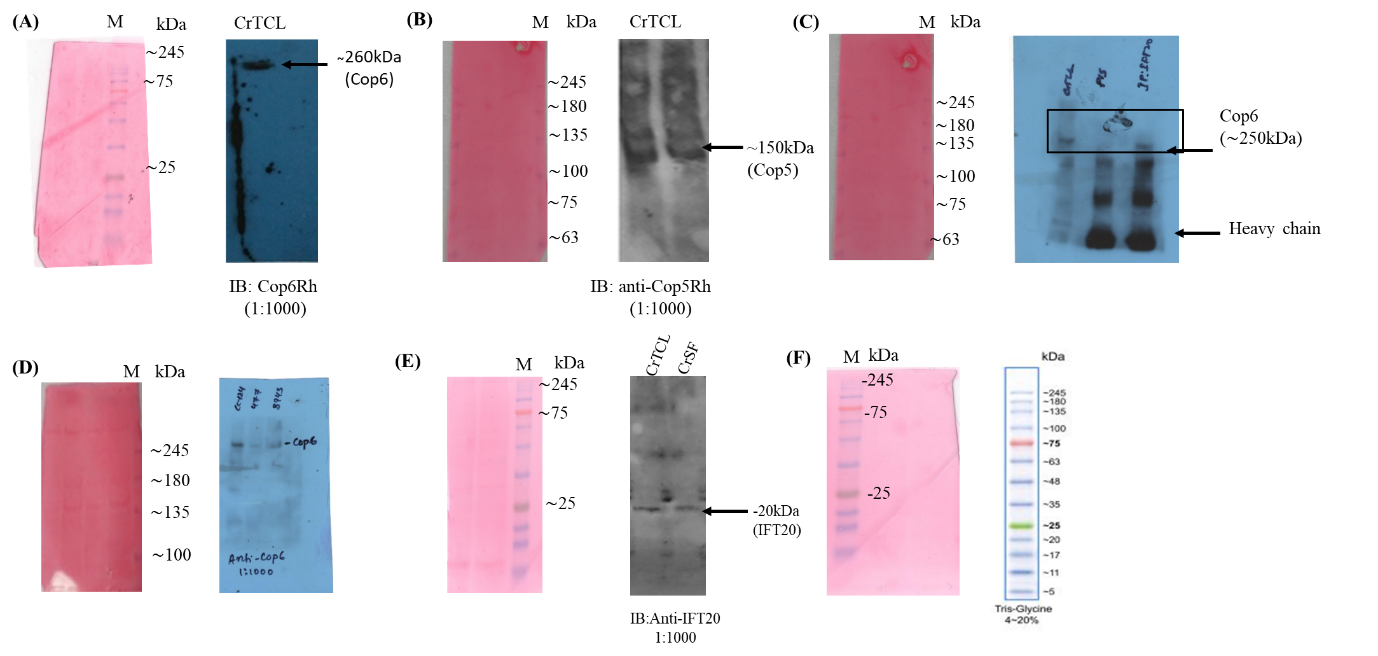
**Appendix figures**

**Appendix Figure 1: Depiction of molecular marker position in the immunoblots.** (A and B) depiction of molecular weight marker in the immunoblot of Cop6 and Cop5, respectively. (C) full blot image of immunoblot with anti-Cop6 after co-immunoprecipitation of Cop6 with Anti-IFT20 (representative of figure 6); band at approximately ∼245kDa in the input lane (CrTCL) and IP: IFT20 is highlighted in a black box. Marker positions are shown alongside in the ponceau staining. (D) Depiction of the molecular weight marker in ponceau S staining for comparing the molecular weight in the immunoblot of Cop6 in wild type, IFT52 and IFT88 mutant strains of *C. reinhardtii.* (E) Depiction of the molecular weight marker in ponceau S staining for comparing the molecular weight in the immunoblot of IFT20 with the cell lysate of *C. reinhardtii*. (F) Representation of molecular weight marker in the immunoblot and compared with the migration patterns of BLUelf (GeneDirex) prestained protein ladder provided by the manufacturer.

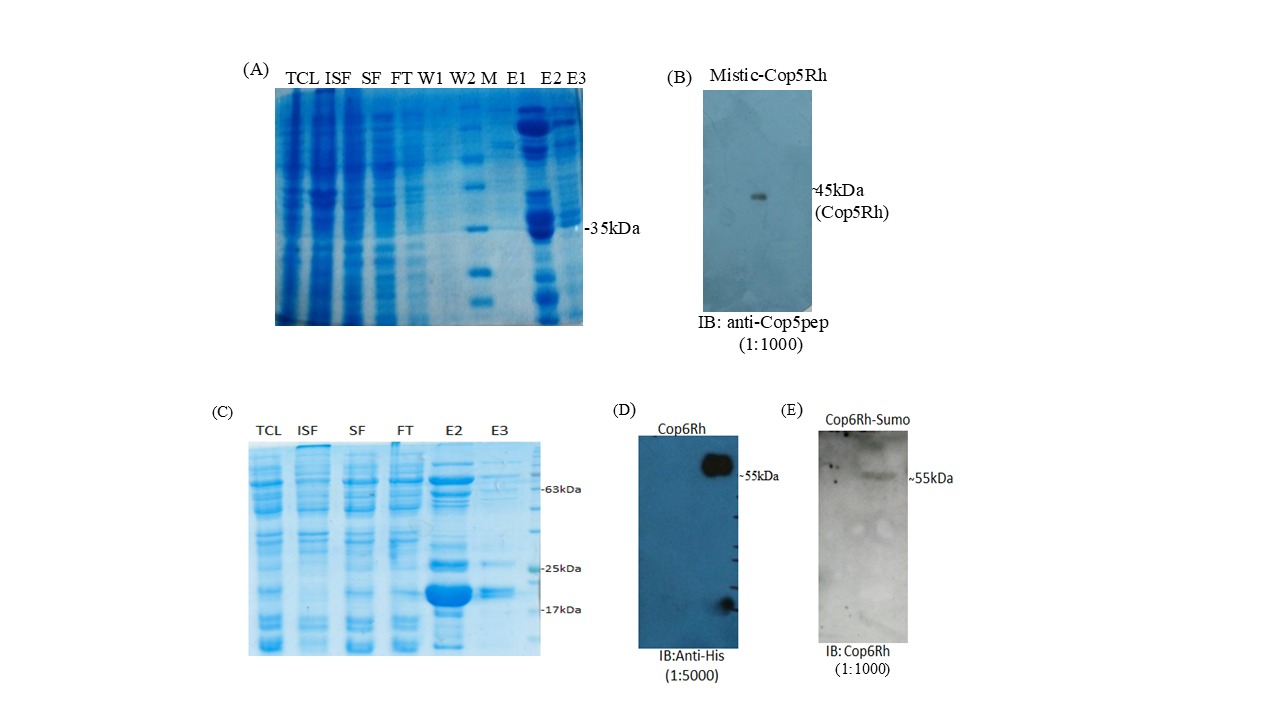

**Appendix figure 2: Determination of the specificity of antibodies raised against Cop5 and Cop6.**

(A) SDS-PAGE of fractions obtained during purification of Cop5 rhodopsin domain; TCL: Total cell lysate, ISF: Insoluble fraction, SF: Soluble fraction, FT: Flow through, W:wash, E: Elution. (B) Immunoblotting of purified rhodopsin domain of Cop5 protein (Elution 1) with anti-Cop5; a band at the expected molecular weight of Cop5 rhodopsin domain (approximately 45kDa) was observed. (C) SDS-PAGE of fractions obtained during purification of Cop6 rhodopsin domain. (D) Immunoblotting of purified rhodopsin domain of Cop6 protein (Elution 1) with anti-His tag; a band at the expected molecular weight of Cop6 rhodopsin domain (approximately 55kDa) was observed. (E) Immunoblotting of purified rhodopsin domain of Cop6 protein (Elution 1) with anti-Cop6; a band was observed at 55kDa, similar to the immunoblot with anti-His.

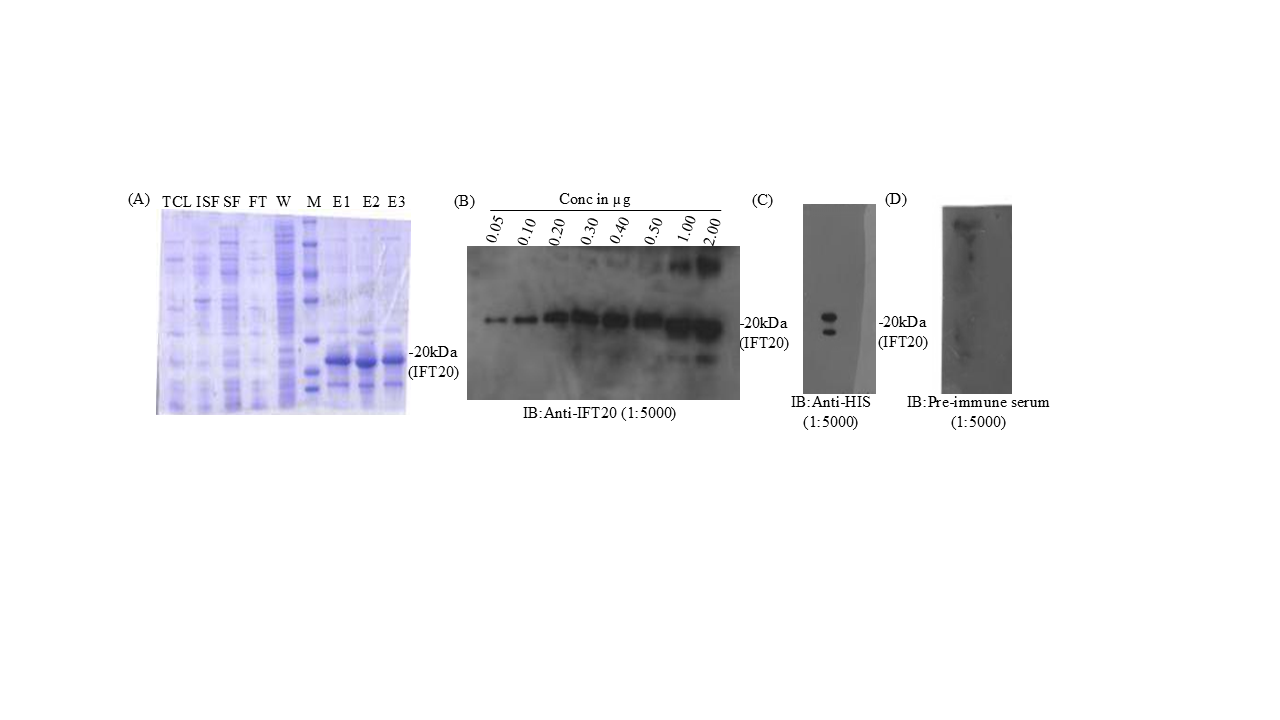
**Appendix figure 3: Determination of the specificity of Anti-IFT20**

(A) SDS-PAGE profile of fractions obtained during the purification of heterologously expressed IFT20. TCL: Total cell lysate, ISF: Insoluble fraction, SF: Soluble fraction, FT: Flow through, W: Wash, E: Elution. (B) The sensitivity of anti-IFT20 was determined by varying the concentration of purified IFT20 and immunoblotted with anti-IFT20. The anti-IFT20 can detect up to 50 ng of purified IFT20 protein. (C) Immunoblotting of purified IFT20 with anti-His. (D) Immunoblotting of purified IFT20 with pre-immune serum.
